# Structural and functional analysis of the Nipah virus polymerase complex

**DOI:** 10.1101/2024.05.29.596445

**Authors:** Side Hu, Heesu Kim, Pan Yang, Zishuo Yu, Barbara Ludeke, Shawna Mobilia, Junhua Pan, Margaret Stratton, Yuemin Bian, Rachel Fearns, Jonathan Abraham

## Abstract

Nipah virus (NiV) is a bat-borne, zoonotic RNA virus that is among the most pathogenic viruses known to humans. The NiV polymerase, which mediates viral genome replication and mRNA transcription, is a drug target. However, NiV polymerase structures were previously unavailable. We determined the cryo-EM structure of the NiV polymerase complex, comprising the large protein (L) and its associated co-factor (P), and performed structural, biophysical, and functional analyses of the NiV polymerase. The complex assembles with a long P tetrameric coiled-coil that is capped by a bundle of ⍺-helices that we show are likely dynamic in solution. Highly conserved zinc-binding modules in the capping domain and a large insert in the RdRp palm domain that is short or absent in most other non-segmented negative strand RNA viruses are critical for replication and transcription. Our findings have the potential to aid in the rational development of drugs to combat NiV infection.

## Introduction

The spillover of RNA viruses from their natural reservoirs into human populations continuously threatens global public health. Nipah virus (NiV) is an enveloped RNA virus of the genus *Henipavirus* in the *Paramyxoviridae* family of the order *Mononegavirales*, the non-segmented negative strand RNA viruses (nsNSVs)^1^. In humans, NiV infection can lead to respiratory illness or encephalitis with a high fatality rate, ranging from 40% to 70%^2^. Since the original identification of NiV in Malaysia in 1998, spillover events have occurred almost annually in Bangladesh, and NiV has also caused outbreaks in India and in the Philipines^3–5^. In 2023, there were NiV outbreaks in Bangladesh and India, and cases have been reported in Bangladesh in 2024.

Although prior NiV outbreaks have been limited in size, NiV may have pandemic potential. This is because infected individuals can be asymptomatic, the virus can be isolated from respiratory secretions, and it can be transmitted from person to person. For example, droplet-based transmission through coughing was observed in the 2018 NiV outbreak in Kerala, India^4^. There is, however, no vaccine or antiviral to combat NiV infection, highlighting a critical gap in public health preparedness. For these reasons, NiV is on the World Health Organization Research & Development Blueprint list of priority diseases for which there is an urgent need for accelerated research and countermeasure development.

The NiV genome is a single strand of negative sense RNA that is transcribed into mRNAs and replicated, to produce progeny genomes, in the cytoplasm of a host cell^6,7^. The viral machinery required for transcription and replication includes the viral genome, packaged by multiple copies of the nucleoprotein (N) into a helical ribonucleoprotein (RNP) complex^8^ and an RNA dependent RNA polymerase. The NiV polymerase is multifunctional, capable of synthesizing subgenomic mRNAs, containing a methylated cap and poly A tail, in addition to synthesizing full-length, encapsidated, replicative RNAs. The NiV polymerase complex comprises the large protein (L) and phosphoprotein (P)^9,10^, with L containing the enzymatic domains necessary for transcription and RNA replication. By analogy with the polymerases of other non-segmented negative strand RNA viruses (nsNSVs) the L protein comprises five domains; the RdRp domain, the capping domain (CAP), the connector domain (CD), the methyltransferase (MTase) domain, and the C-terminal domain (CTD)^11–13^. The P protein serves an adaptor that allows the polymerase to associate both with the RNP template and with soluble N protein that is required to concurrently encapsidate newly synthesized replicative RNA. P contains an intrinsically disordered N-terminal domain (NTD), a central oligomerization domain (OD), and a C-terminal X domain (XD). The C-terminal domain of P binds the RNP template, and the N-terminal domain binds soluble N protein for encapsidation^14–17^.

Despite the public health relevance of NiV as a human pathogen, no NiV L–P structures are available to help guide rational drug development efforts. This is an important gap in knowledge because viral polymerases are critical drug targets against many RNA and DNA viruses. For example, the nucleotide prodrug remdesivir, which targets NiV L activity, has demonstrated preclinical efficacy against lethal NiV infection in non-human primates^18,19^. Although structures of other nsNSV polymerases are available, rational drug design efforts against the NiV polymerase complex has been limited by lack of an understanding of its unique and/or conserved features that may serve as suitable drug targets.

Here, we determined the 2.3 Å cryo-EM structure of NiV L–P complex, which reveals how tetrameric P interacts with L. Cryo-EM 3D-variability analysis paired with molecular dynamics simulations suggest that the distal end of the long tetrameric P coiled-coil is highly flexible. Structure-function analysis of the L–P complex using a NiV minigenome assay identifies both conserved and unique features of the polymerase that are required for transcription and genome replication, providing information that has the potential to aid rational design of antiviral molecules against NiV.

## Results

### Characterization of NiV L–Pcomplexes

We co-expressed full-length NiV L (residues 1–2244) and full-length P (residues 1–709) proteins in insect cells (Figure 1A). We further purified the L–P complex using affinity chromatography followed by size exclusion chromatography and SDS-PAGE analysis (Figure S1A). The identities of the L and P bands were confirmed by LC-MS/MS, which revealed high coverage for the full-length proteins (Figure S2), and analysis of samples by mass photometry revealed a major peak of 559 kDa, which likely represents L bound to four copies of P (i.e., an L– P_4_ complex) (Figure 1B). We found the purified NiV L–P complex to be active in an in vitro RNA synthesis assay using an oligonucleotide RNA template containing the NiV promoter sequence and nucleotide triphosphates (NTPs), including [*α*^32^P] GTP as a radioactive tracer (Figures 1C and 1D). This analysis showed that the L–P complex was functional.

**Figure 1.**
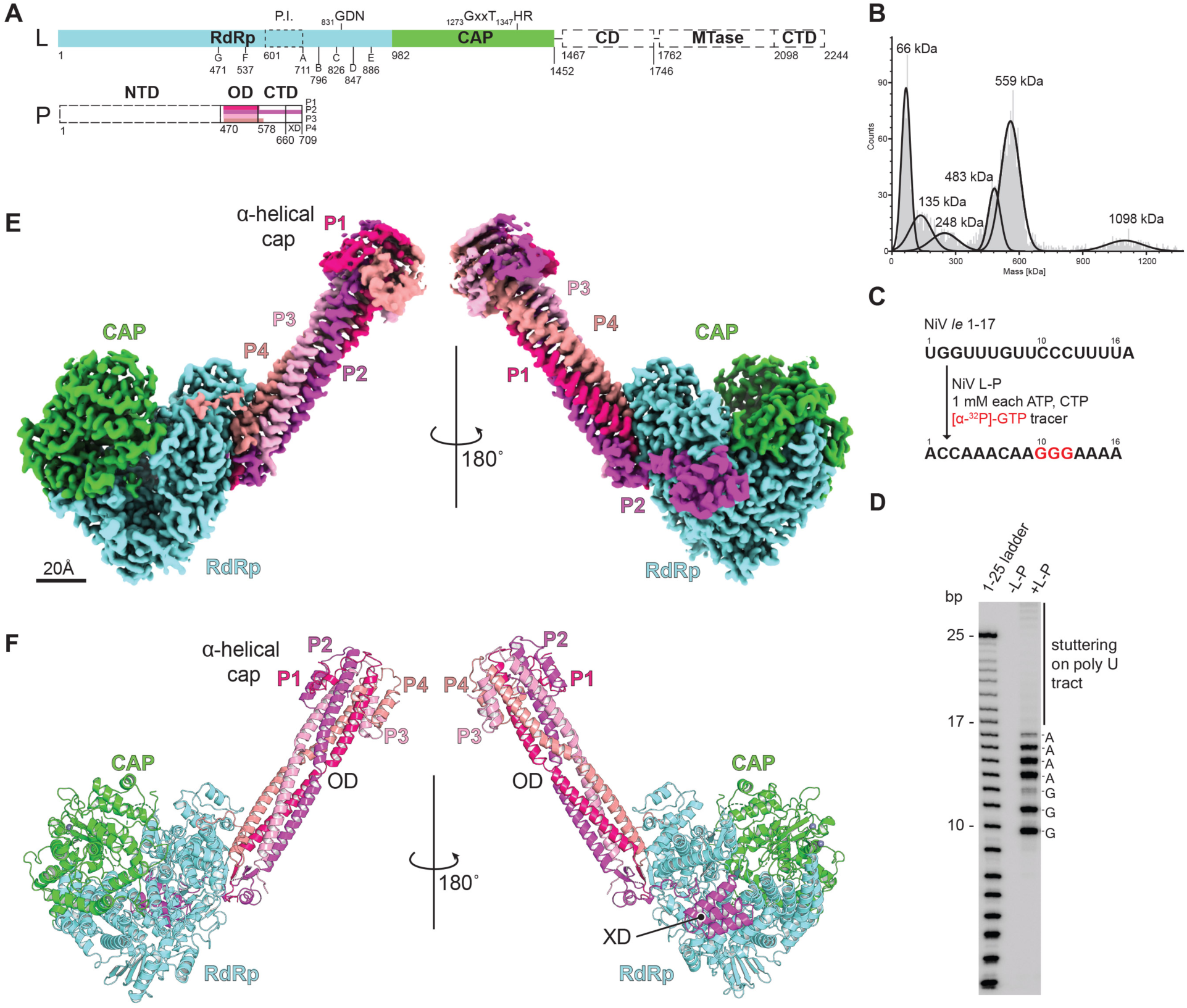
Structure of the NiV L–P complex. (A) Schematic diagram of the domain organization of the NiV L and P proteins. L contains five domains which are the RdRP, capping (CAP), connector (CD), methyltransferase (MTase), and C-terminal (CTD) domains. Only the RdRp domain and capping domains, which form the polymerase core, could be resolved; CD, MTase, and CTD domains (dashed boxes) had no cryo-EM density and could not be modelled, suggesting that they are flexible with respect to the L core. P contains three domains, which are the N-terminal domain (NTD), the oligomerization domain (OD), and the C-terminal domain; the X domain (XD) is within the CTD. The regions of tetrameric P protein that could be resolved vary in length depending on the protomer (P1–P4). The P NTD of each protomer (dashed box) could not be resolved likely due to flexibility. P NTD, OD, CTD, and XD boundaries are labeled based on established boundaries^14^. (B) Mass photometry result of NiV L–P complex. Based on expected molecular masses, the 559 kDa peak likely represents the L–P_4_ complex, the 483 kDa complex would be consistent with an L–P_3_ complex, the 248 kDa peak may correspond to the L protein alone, and the 1098 kDa could represent dimers of L–P_4_. The 135 kDa and 66 kDa peaks may represent degradation products or contaminants. This experiment was performed twice, with representative data shown. (C) Flow diagram of the RNA synthesis assay. The template is the leader sequence of NiV, nucleotides 1–17. The GTP tracer and its incorporation in the product are in red. (D) RNA synthesis activity assay with radioactive products migrated on a denaturing polyacrylamide gel. nt; nucleotide. This experiment was performed a total of four times with two different L–P preparations, with representative data shown. (E) The cryo-EM density map of NiV L–P complex with L domains and P protomers colored as indicated. Two views are shown. (F) Ribbon diagram of the NiV L–P complex with L domains and P protomers colored as indicated

### Cryo-EM structure of NiV L–P complex

We next used single-particle cryo-electron microscopy (cryo-EM) to determine the structure of the NiV L–P complex to a global resolution of 2.3 Å (Figures S1B–D). The L–P complex is shaped like a tobacco pipe, with the stem formed by the tetrameric P OD (Figures 1E and 1F). We observed interpretable density for L residues 5–1462, encompassing the RdRp and CAP domains, and for the P tetramer (P1–P4), of which we could observe different lengths for individual protomers (P1: residues 479–584, P2: 479–708, P3: 477–579, and P4 479–596) (Figure 1A). We did not observe density for the L CD, CTD, and MTase domains, despite using the full-length L protein in our preparations as confirmed by LC-MS/MS and based on the mass of assembled L–P_4_ complexes detected by mass photometry (Figures 1B and S2). Therefore, like in structures of full-length L proteins of respiratory syncytial virus (RSV) and human metapneumovirus (HMPV) (*Pneumoviridae*), the three L C-terminal globular domains in NiV L are probably too flexible relative to the RdRp-CAP polymerase core to be visualized^20–25^. For the domains that could be resolved (RdRp and CAP), alignment between NiV L and other nsNSV L based on the Cα ranged from 1.7 to 5.0 Å (Supplementary Table 2), indicating structural conservation, consistent with the generally similar transcription and replication mechanisms of nsNSVs.

In cryo-EM maps, the N-terminal region of the NiV P tetramer is capped by a mushroom-shaped density (Figures 1E and S1B). Focused refinement centered on this region nonetheless allowed us to resolve individual ⍺-helices, allowing for unambiguous modelling of the region based on an available X-ray crystal structure of the P OD domain^16^. The ⍺-helices form a bundle that folds back onto the outer aspects of the tetrameric coiled-coil core (Figures 1F), which is a feature that has also been observed in X-ray crystal structures of the NiV and Sendai virus (paramyxovirus) phosphoproteins^16,26,27^.

### Structural features of the NiV L protein

The NiV L protein RdRp domain has a conventional right-hand “fingers-palm-thumb” organization containing seven motifs involved in catalysis (A–G), like other nsNSV RdRp domains (Figures 2A and 2B). Motifs A–E are in the palm while motifs F and G are in the fingers. Motif C contains the active site residues GDNE (residues 831–834), which are part of the active site. The terminal glutamate in this motif is replaced by a glutamine in the L proteins of almost all nsNSV members we examined and is only a glutamate in a few species including NiV and Hendra virus (Figure S3). However, glutamate and glutamine were previously shown to be interchangeable at this position in NiV L without a detectable effect on enzyme activity^28^.

**Figure 2.**
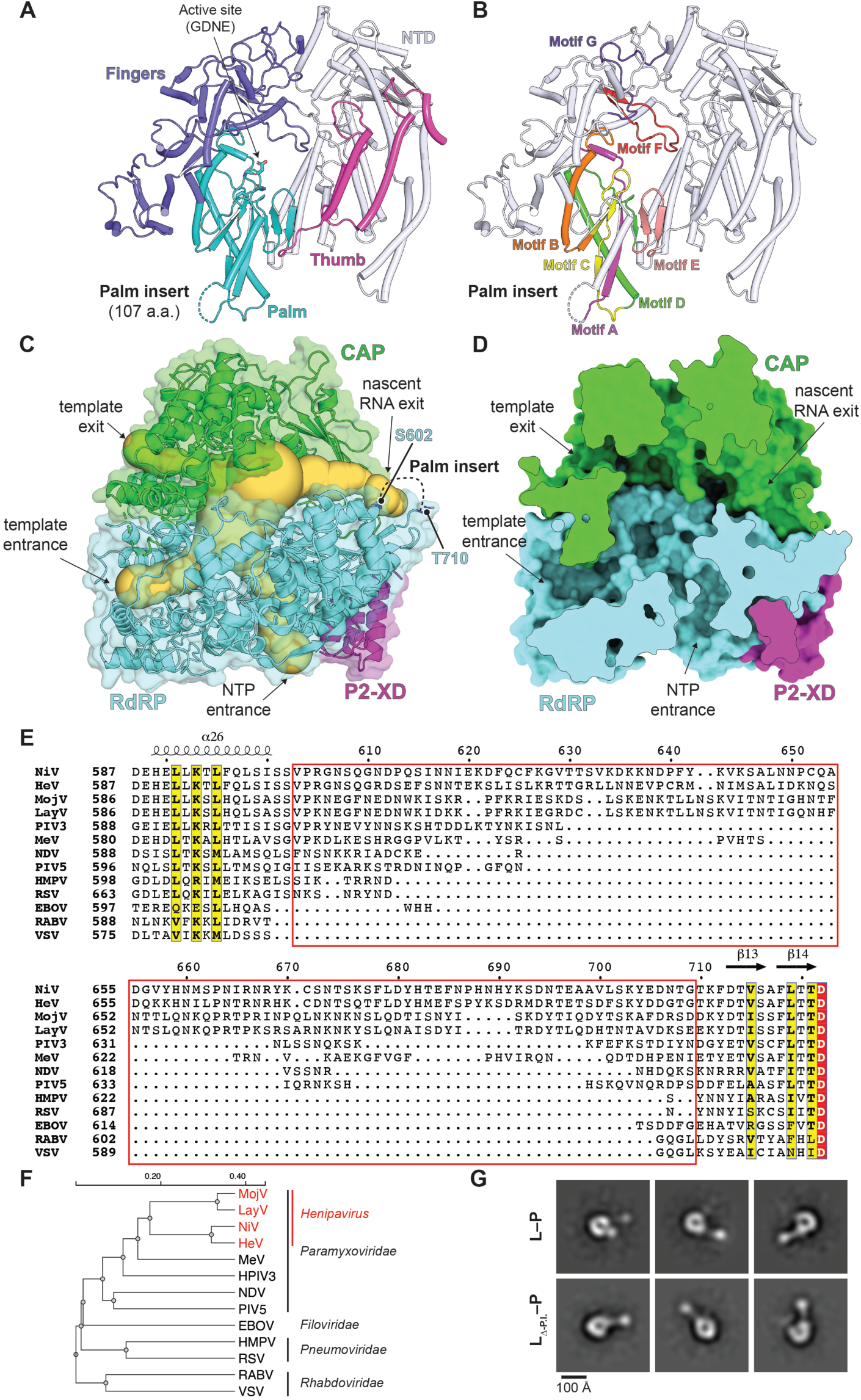
Molecular features of the NiV L polymerase core. (A) Ribbon diagram of the NiV L RdRp domain. The fingers, palm, and thumb domains are shown and colored as indicated. Palm active site residues (GDNE) are shown as sticks. The location of the large palm insert is indicated. (B) The same view as in *A*, with catalytic motifs A–G colored as indicated. (C) Structure of NiV L– P complex shown as a ribbon diagram with transparent surface with channels shown as spheres (calculated by CAVER Web^44^). The template entrance, template exit, NTP entrance, and nascent RNA exit channels are indicated. Residues S602 and T710, near the two ends of the palm insert, are located near the nascent RNA exit channel. (D) Clipped surface of the NiV L–P complex showing the RNA synthesis channels in the same view as shown in *C*. (E) Sequence alignment of the region of the palm insert for the different nonsegmented negative-strand RNA virus (nsNSV) L protein sequences. The NiV palm insert (amino acid residues 603–709) is boxed in red. NiV, Nipah virus; HeV, Hendra virus; MojV, Mojiang virus; LayV, Langya virus; PIV3, human parainfluenza virus 3; MeV, measles virus; NDV, Newcastle disease virus; PIV5, parainfluenza virus 5; HMPV, human metapneumovirus; RSV, respiratory syncytial virus; EBOV, Ebola virus; RABV, rabies virus; VSV, vesicular stomatitis virus. See Methods for GenBank accession numbers. (F) Phylogenetic tree of the indicated nsNSVs based on L amino acid sequences generated using Clustal Omega^45^. Genera and families of each virus are indicated. See Table S3 for additional information. (G) Negative stain EM 2D classes of purified NiV L–P complexes for wild-type (WT) L or an L mutant in which residues 604–704 of L (L_Δ-P.I._). P.I. stands for palm insert. See Figure S4F–H for additional information.

On the assembled complex, there are channels for nucleotide triphosphate (NTP) entrance, template entry, template exit, and nascent RNA exit (Figures 2C and 2D). The template entrance channel can be identified by analogy to the Ebola virus (EBOV) and RSV L proteins, which have recently been determined in complex with their cognate promoter RNAs^20,29^. This channel involves surfaces of the RdRp and capping domains (Figure 2D). The putative template exit channel, identified by analogy with other viral RdRps, is within the capping domain (Figures 2C and 2D)^11^. The putative nascent RNA exit channel is formed by the RdRp domain and the CAP domain on the other side of L relative to the template entrance channel.

An interesting distinction between the NiV RdRp domain and that of most other nsNSV polymerases lies in the loop found between residues 600–713 (NiV L numbering), which is located immediately N-terminal to motif A in the palm domain (Figures 2A and 2B). We did not observe cryo-EM density for residues 603–709 within this loop. Amino acid sequence alignments of L proteins revealed that this region is highly variable between nsNSVs (Figures 2E and S4)^30,31^. Compared to other nsNSVs, including most other paramyxoviruses, henipaviruses have additional sequence at this site, which we refer to as the palm insert, which ranges from ∼100 to more than 200 amino acid residues in length, and is remarkably longer than the analogous sequences in other nsNSVs. For example, the shortest analogous loop in the L proteins of vesicular stomatitis virus (VSV) and rabies virus (RABV) (*Rhabdoviridae*) and is only eight residues long (Figure 2E). Other than the large size and general features of being rich in lysine and asparagine residues, there is substantial divergence in the sequence of the loop even among henipaviruses. However, the loop is mostly conserved among closely related henipaviruses when they are examined as pairs (e.g., NiV and HeV, MojV and LayV), with Cedar virus having a particularly long insert (>200 residues) (Figures 2F and S4A–E).

Examination of the palm insert region in the cryo-EM maps showed that amino acids 603– 709 are not visualized, suggesting a high degree of flexibility. However, the boundary residues that we could observe for the palm insert, S602 and T710, are relatively close to each other (only ∼10 Å apart) in the structure and are located near the putative nascent RNA exit channel (Figure 2C). The proximity of the N- and C-terminal regions of the loop and disorder in cryo-EM maps suggest that the insert may form an appendage that does not otherwise contribute to the fold of the RdRp domain. To test this hypothesis, we generated a recombinant form of NiV L that lacks residues 604–704 (NiV L_Δ-P.I._) and co-expressed this mutant L protein with P in insect cells. Purified particles when examined by negative stain electron microscopy had the overall similar tobacco pipe shape as did particles for the wild-type NiV L–P complex (Figures 2G and S4F–H). These findings confirmed that the unique palm insert does not contribute to the fold of the RdRp domain.

### Capping domain

The CAP domain of nsNSV polymerases is multifunctional, playing roles in RNA synthesis initiation and regulation of elongation, in addition to catalyzing the polyribonucleotidyltransferase (PRNTase) step of cap addition^11–13^. For VSV, the CAP domain has also been demonstrated to interact with the N protein of the template RNP^32^. Key features that have been identified in other nsNSV polymerases include zinc finger domains of unknown function^29,33–38^, a priming loop, thought to be involved in stabilizing the RNA synthesis initiation complex, and an intrusion loop, which contains the catalytic HR motif required for PRNTase activity^11–13^. The priming and intrusion loops have been captured in multiple distinct conformations for different nsNSV polymerases, with each thought to represent distinct functional states (Figure S5). In the cryo-EM structure of the NiV L–P complex, most of the priming loop and intrusion loop are not visible in cryo-EM maps (amino acids 1266–1289 and 1342–1362, respectively), likely due to their flexible nature (Figure 3A). This state captured with NiV L–P is most similar to Ebola L–VP35 complexes visualized in the absence of RNA, in which most of the priming loop and intrusion loops were disordered, with these loops only becoming ordered in the presence of an RNA template-primer duple (Figure 3C–E)^29,38^.

**Figure 3.**
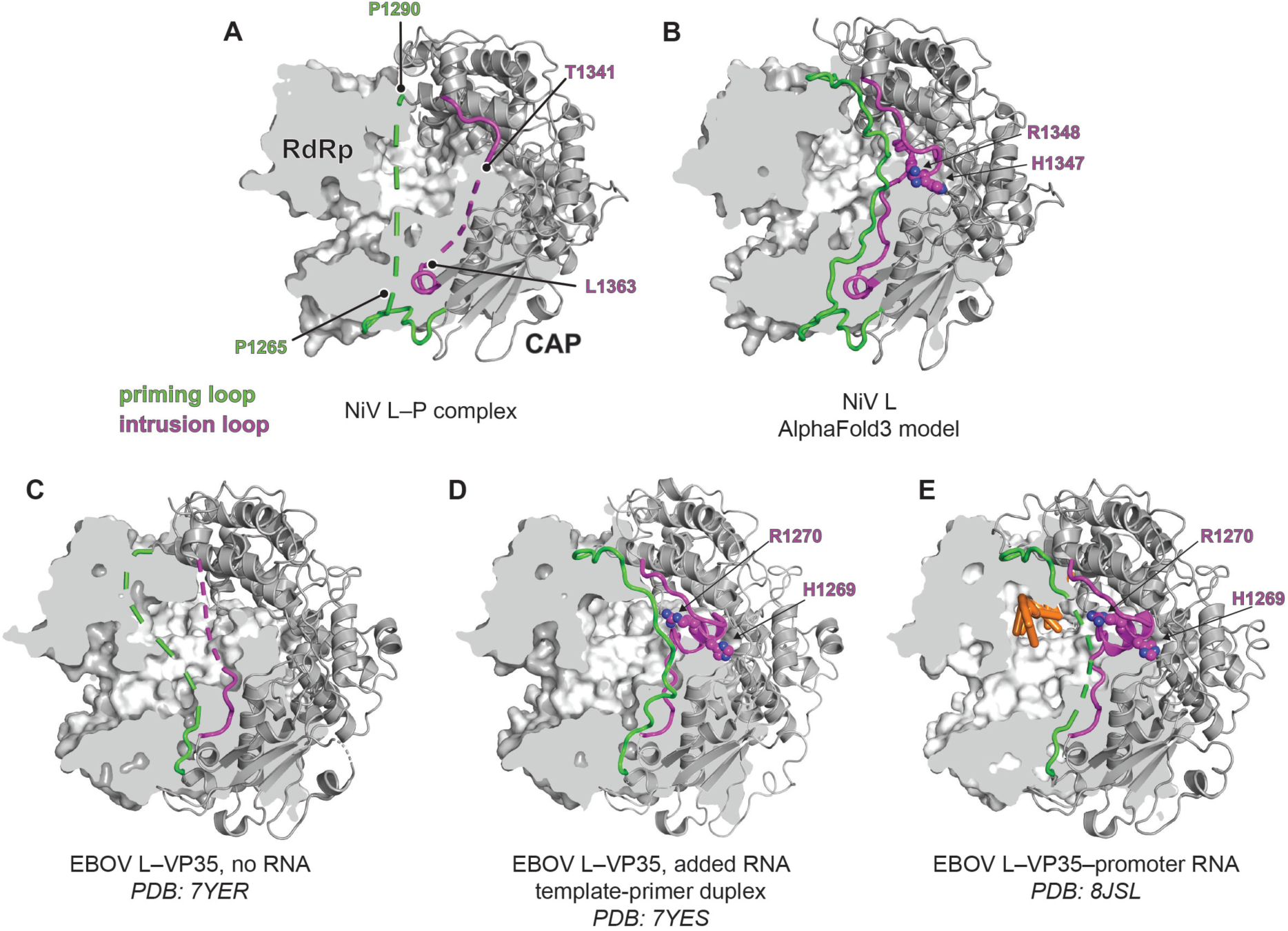
Priming and intrusion loop conformations in the NiV L capping domain. (A) The NiV L protein RdRp domain is shown in surface representation with a partially clipped surface, and the CAP domain is shown as a ribbon diagram. The priming (green) and intrusion (purple) loops are disordered in the cryo-EM structure. Boundary residues are indicated. The disordered portions are shown as dashed lines. (B) Predictions of the NiV L protein RdRp and CAP domains derived from an AlphaFold3^46^ model generated in the absence of nucleic acids, represented as shown for the cryo-EM structure in *A*. Intrusion loop HR residues (H1347 and R1348) are shown as spheres. (C–E) Cryo-EM structure of EBOV L-VP35 without addition of RNA (PDB: 7YER)^38^ (C), with the addition of an RNA template-primer duplex with (with no RNA density observed in maps; PDB: 7YES)^38^ (D), and with promoter RNA and nucleic acids observed (PDB: 8JSL)^29^ (E). Nucleic acids are shown as orange sticks. Intrusion loop HR residues are also shown as sticks. For the EBOV L–VP35 complex, addition of an RNA template-primer duplex was required to visualize the priming and intrusion loops.

### Overall structure of the NiV P protein

In the assembled complex, as in structures of P proteins bound to the L proteins of other paramyxoviruses^33,37,39^, most of the N-terminal region of P remains disordered (Figure 1A). The C-terminal regions of P that are visualized in each of the four P protomers fold onto L by forming several extended arms, with one of the protomers (designated P2 here) providing contacts through XD domain, which forms a bundle of three ⍺-helices (⍺-1, ⍺-2, and ⍺-3) (Figure 4). Of the P4 XD helices, ⍺-1 and ⍺-3 interact with L (Figure 4D). The P “fold upon binding” interaction mode has been extensively described also for other nNSVs^21,22,24,33,37,39^. In the L–P complex, P buries approximately 3478 Å^2^ of surface area on L. Only three of the four protomers (P1, P2, and P4) contact the L protein, with the extended arms of P1 and P4 forming an extensive network of polar interactions with the RdRp (Figures 4B–E). As part of this network, P1 interacts with the RdRp through β-strand augmentation, which is further stabilized by a salt-bridge formed by the P2 residue R600, whose side chain reaches across the added β-strand to interact with RdRp residue D384 (Figure 4B). The P2 polypeptide chain makes additional polar contacts as it courses towards the XD domain (Figure 4C). An extension in P4 interacts with the RdRp domain on the opposite face that binds P2 XD (Figure 4E). As part of this extension, the side chains of P4 residues K583, K587, and K589 make polar interactions that anchor the loop onto the RdRp domain (Figure 4E).

**Figure 4.**
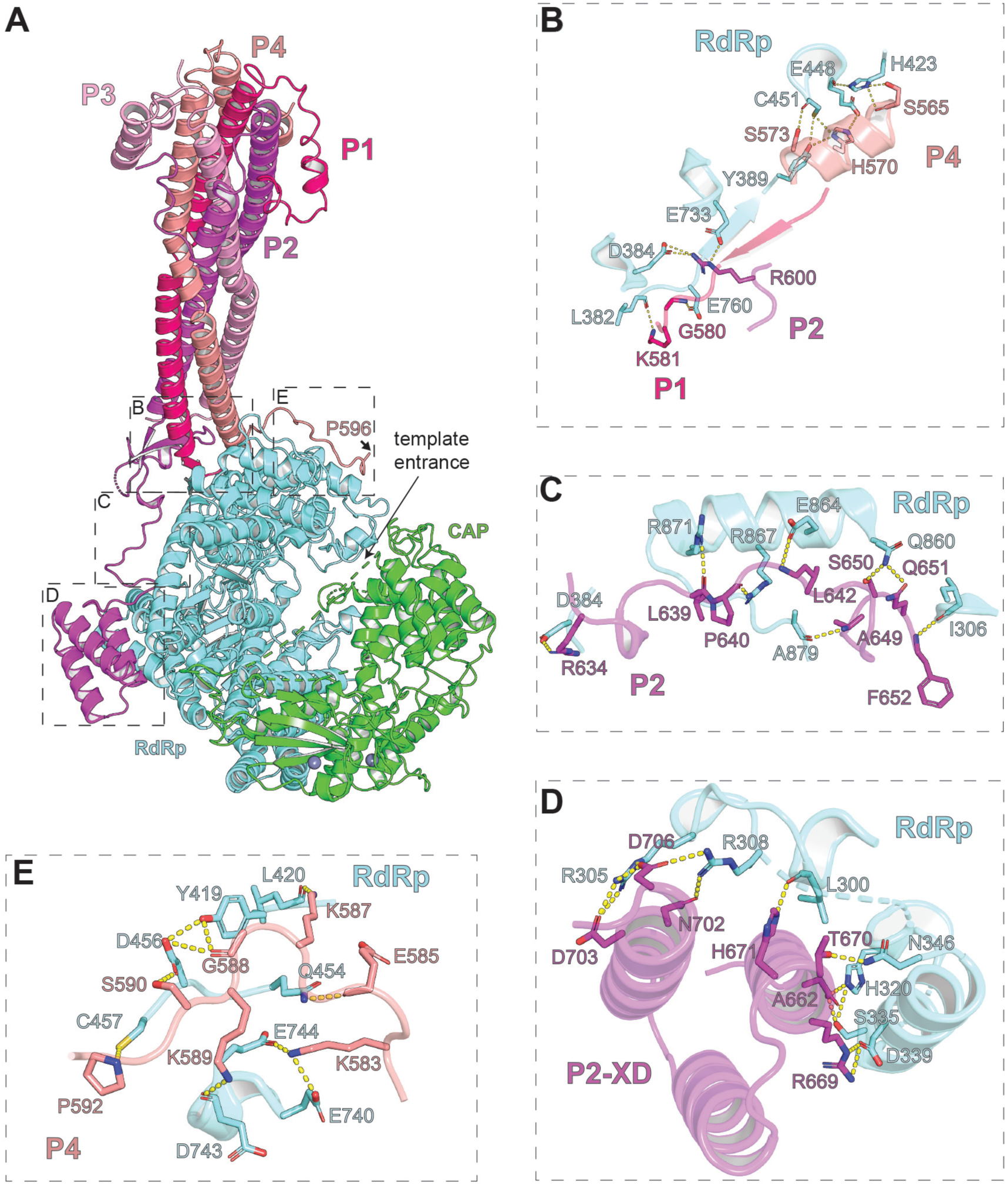
Interactions between NiV P and L RdRp. (A) Ribbon diagram of the NiV L–P complex. The interaction interfaces between tetrameric P and NiV L occurs over four main regions (B–E). (B) Interactions between the P1, P2, and P4 and the RdRp domain. Interacting side chains and main chain atoms are shown sticks. (C) Interactions between the P2 linker, which connects the P2 OD with P2 XD, and the L RdRp are shown. (D) Interactions between the P2 XD ⍺-helices and the L RdRp. (E) Interaction the P4 peptide extension and the L RdRp.

### Dynamics of L-associated phosphoprotein

Despite co-expressing and purifying preparations of full-length P and L, as in other structure of L–P complexes, we were only able to resolve about the C-terminal one third of P. Our structure nonetheless contains the longest P segment visualized in paramyxovirus L–P complexes, particularly when the folded back helices of the alpha-helical cap structure are considered, as the two bent helices add an additional ∼50 Å to the 100 Å coiled-core (Figure S9).

We hypothesized that the extensive network of polar interactions P makes with L would likely be malleable, allowing the pose of the P OD stem to vary with respect to the polymerase core. Consistent with such motion, the distal (N-terminal tip) of P had the lowest local resolution, with features of the ⍺-helical cap structures suggestive of movement (Figures 1E and S1D). We thus conducted 3D variability analysis^40^ on cryo-EM datasets of assembled complexes, which revealed a swiveling motion of the P OD with respect to the polymerase core (Figure 5D and Movie S1). As an additional method to characterize P movements with respect to the L polymerase core, we performed 100 nanosecond molecular dynamics simulations using the cryo-EM structure as a starting point (Figure 5A–C and Movies S2 and S3). Molecular dynamics analysis revealed that mean root square fluctuations (RMSF) were much higher for each of the for P protomers than they were for the polymerase core (RdRp/CAP), with the highest fluctuations observed in the N-terminal portions of the ⍺-helical cap structure (Figure 5A–C and Movies S2 and S3). Structural fluctuations were also notably high for the C-terminal region of P4, which forms the P4 extension noted above and wraps around the RdRp domain near the template entrance channel (Figure 4E and Movies S2 and S3).

**Figure 5.**
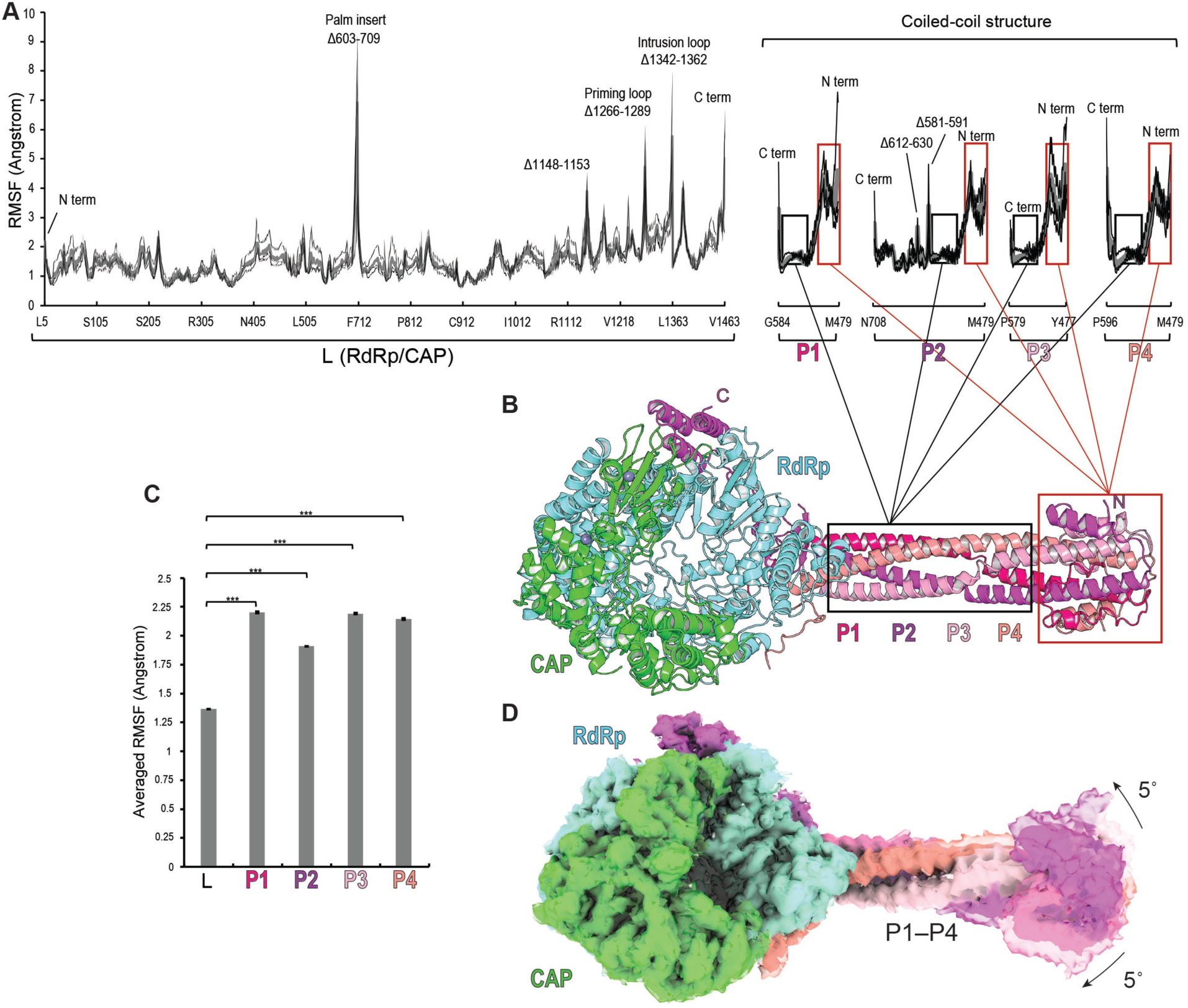
Tetrameric P is flexible when assembled onto the L polymerase core. (A–B) Per-residue root-mean-square-fluctuation (RMSF) from 100 ns molecular dynamics (MD) simulations of the protein complex (A) with ribbon diagram of the cryo-EM structure of the NiV L–P complex provided as a reference for regions of P with increased flexibility. Three thin lines represent individual MD runs. A thick line represents averaged RMSF values. For the RdRp/CAP portion of the plot, regions noted as “Δ” are stretches of residues that had no observable or interpretable density in cryo-EM maps and therefore were not modeled in atomic coordinates; high RMSF fluctuations in these regions is likely due to interruption of the polypeptide chain. (C) Mean RMSF values calculated during MD simulations for L(RdRp/CAP), P1, P2, P3, and P4 are plotted. The comparison of mean RMSF values between two groups was performed using an unpaired, two-tailed Student’s t-test. ***p<0.001. (D) 3D variability analysis^40^ on NiV L–P dataset reveals a swiveling motion of the P OD with respect to the L core. See Movie S1.

### Functional analysis of key features of the polymerase

As noted above, the NiV polymerase is responsible for both transcription and genome replication, distinct processes that each involve a different sequence of events. During transcription, the polymerase starts from a promoter in the *leader* (*le*) region at the 3’ end of the genome and moves along the template RNP, responding to cis-acting *gene start* and *gene end* sequences to produce a series of subgenomic mRNAs each with a 5’ methylguanosine cap and 3’ polyadenylate tail. During replication, the *gene start* and *gene end* signals are not recognized and the polymerase produces a full length, uncapped, non-polyadenylated antigenome, which is concurrently encapsidated by N protein. The encapsidated antigenome in turn serves as a template for synthesis of encapsidated genome RNA. We sought to determine if key features identified in the cryo-EM structure of the NiV L–P complex are required for transcription and/ or RNA replication.

We focused our analysis on the following features: (1) the RdRp palm insert (Figure 2); (2) the HR motif in the intrusion loop of the CAP domain (Figures 3A and 3B); (3) The P4 extension that contacts the RdRp (Figures 4E and 6A), (3) and the two zinc finger motifs in the capping domain (Figures 6B and 6C). To determine the importance of these features, we examined NiV L mutants using a cell-based minigenome system, in which NiV minigenome and NiV N, P, and L protein expression is driven by intracellular expression of T7 RNA polymerase (Figure 6D). We used a dicistronic minigenome in which the first gene contains *chloramphenicol acetyltransferase* (CAT) reporter gene sequence, and the second gene contains a *Renilla luciferase* reporter gene. In this assay system, the minigenome can undergo multiple cycles of RNA replication and can be transcribed into CAT and Renilla luciferase mRNAs. Thus, mRNA levels and Renilla luciferase activity correlates with the combined efficiencies of mRNA transcription and replication. For all assays, the parental L plasmid we used for mutational analysis expressed an L protein containing a C-terminal strep tag, to allow monitoring of mutant L protein expression. Comparison of untagged wild-type L and strep-tagged wild-type L in a minigenome luciferase assay showed that the tag had minimal effect on polymerase activity, with luciferase expression from strep-tagged L being at 85% of untagged L levels (data not shown). As a negative control we generated an L plasmid that contained a D832A substitution in the RdRp palm active site GDNE catalytic motif (Figure 2A), which should yield an inactive polymerase.

**Figure 6.**
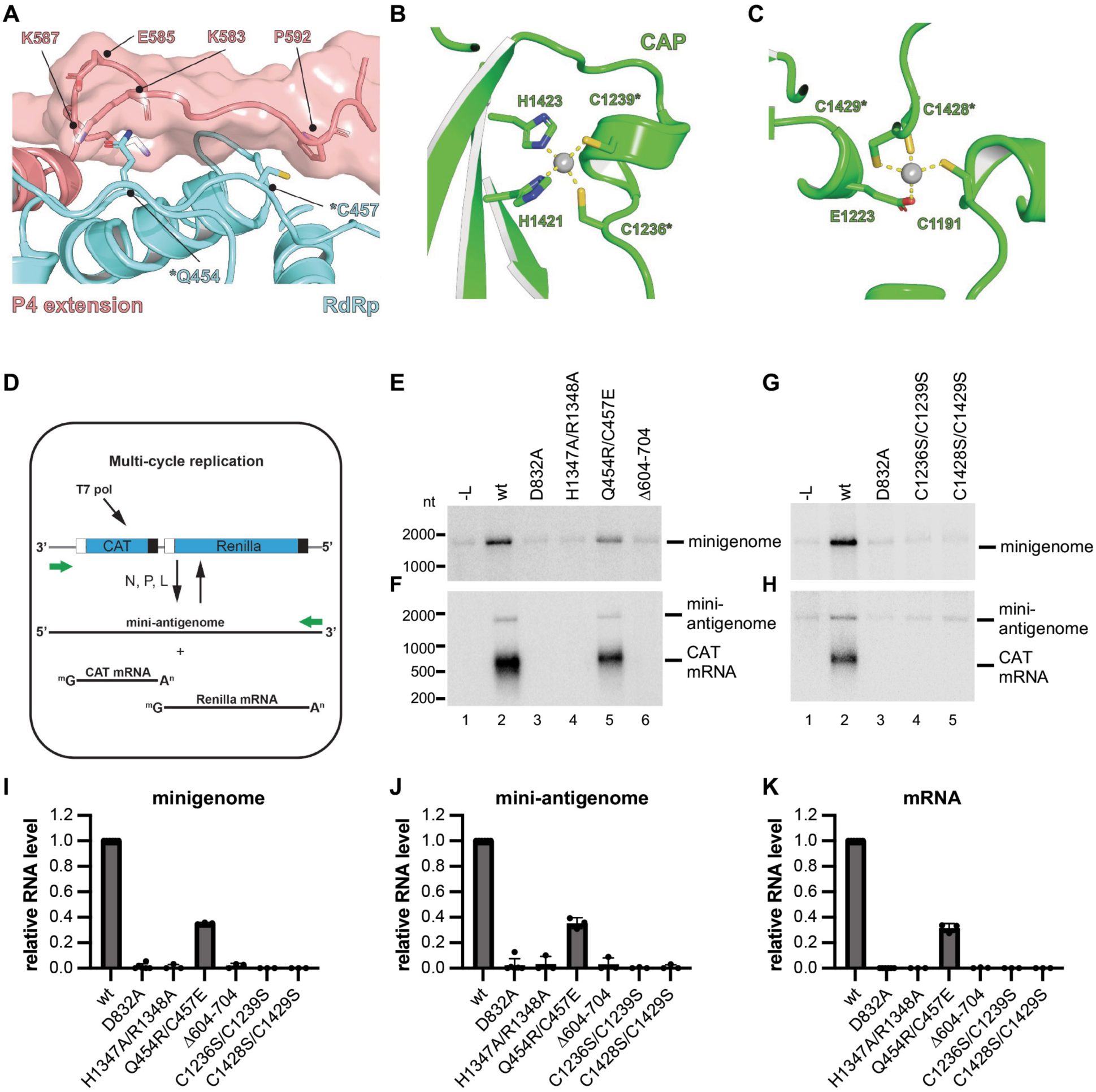
Analysis of NiV L mutants using a multicycle-replication minigenome system. (A) Zoom in view of the P4 extension contact sites on the RdRp. A segment of the P4 extension is shown as a ribbon diagram and also in semi-transparent surface representation. L protein RdRp residues Q454 and C457, indicated by asterisks, are buried during interactions with P4 and were chosen for mutational analysis to disrupt the RdRp interaction with P4. (B–C) Ribbon diagram of the two zinc-binding motifs found in the CAP domain. Cysteines indicated with an asterisk were selected for mutational analysis. See Figure S10 for additional information. (D) Schematic diagram illustrating the structure of the minigenome and the products that are generated during RNA replication and mRNA transcription. White and black boxes indicate gene start and gene end signals, respectively. Green arrows indicate promoters. (E–H) Northern blot analysis of RNAs generated in the minigenome system. Panels E and G show negative sense minigenome template RNA generated by T7 RNA polymerase and amplified by NiV polymerase. Panels F and H show Northern blot analysis of positive sense mini-antigenome RNA and CAT mRNA generated by the NiV polymerase. (I) Quantification of levels of minigenome from replicates of the Northern blots shown in E and G. (J–K) Quantification of levels of mini-antigenome (J) and mRNA (K) from replicates of the Northern blots shown in F and H.

Mutations were introduced into L to delete the palm insert in the RdRp domain (Δ604– 704), to substitute the HR motif in the intrusion loop (H1347A/R1348A), to disrupt the interface with the P4 extension (Q454R/C457E), and to individually disrupt the two zinc finger motifs (C1236S/C1239S and C1248S/C1249S). Transcription and replication were reconstituted in T7 RNA polymerase-expression BSR-T7 cells^41^ (Figure 6D). Western blot analysis confirmed that all mutant L proteins were efficiently expressed, although typically a cleavage product of ∼195 kDa was also detected (Figure S6). This product is the appropriate size to represent the C-terminal portion of L following cleavage at the palm insert region, and consistent with this, the Δ604–704 mutant did not show evidence of this cleavage product. Both zinc finger mutants were also consistently less prone to this cleavage. The Cys2His2 zinc mutant (C1236S/C1239S) was consistently expressed at a lower level than wild-type L and the other mutants (∼ 45% of wild-type levels), suggesting that this zinc finger helps to maintain structural integrity of the L protein. Nonetheless, sufficient levels of this protein were expressed for functional analysis. Northern blot analysis using a CAT-specific probe showed that HR substitutions, both sets of zinc finger mutations, and the Δ604–704 mutation reduced genome and mini-antigenome RNAs to background levels, demonstrating that mutation of these residues abolishes RNA replication (Figures 6E–K). The L Q454R/C457E mutation also impaired RNA replication, reducing genome and mini-antigenome levels to ∼35% of wild-type levels. The Northern blot analysis also showed a complete inhibition of mRNA accumulation in reactions with HR, zinc finger mutants, and the Δ604–704 mutants, and a reduction of mRNA levels in the case of the L Q454R/C457E mutant (Figures 6E–K). The reductions in mRNA levels were mirrored by reductions in luciferase activity (Figure S7A).

The results described above show that the mutations inhibit RNA replication, but the reduction in mRNA and luciferase levels could either be because of inhibition of transcription per se, or because of the reduction in minigenome template levels due to replication inhibition. To determine if the mutations inhibit transcription in addition to replication, we performed additional analyses using a single step minigenome assay that allows uncoupling of these processes. In this assay, the minigenome contains mutations that ablates the promoter at the 3’ end of the mini-antigenome and thus limit the minigenome to the mini-antigenome synthesis step of replication. This allows the effects of mutations on transcription to be assessed independently of any effect that they have on RNA replication (Figures 7A and S8). Analysis of CAT mRNA levels from the single step minigenome showed that the LΔ604–704, HR and zinc finger mutants generated no detectable product, whereas the L Q454R/C457E mutant generated CAT mRNA at ∼ 45% of wild-type levels (Figures 7B–F). This decrease in transcription was validated by analysis of Renilla luciferase levels (Figures S7B). Thus, each of the mutations inhibit transcription in addition to RNA replication, indicating that the L protein features that were mutated are fundamental to polymerase activity.

**Figure 7.**
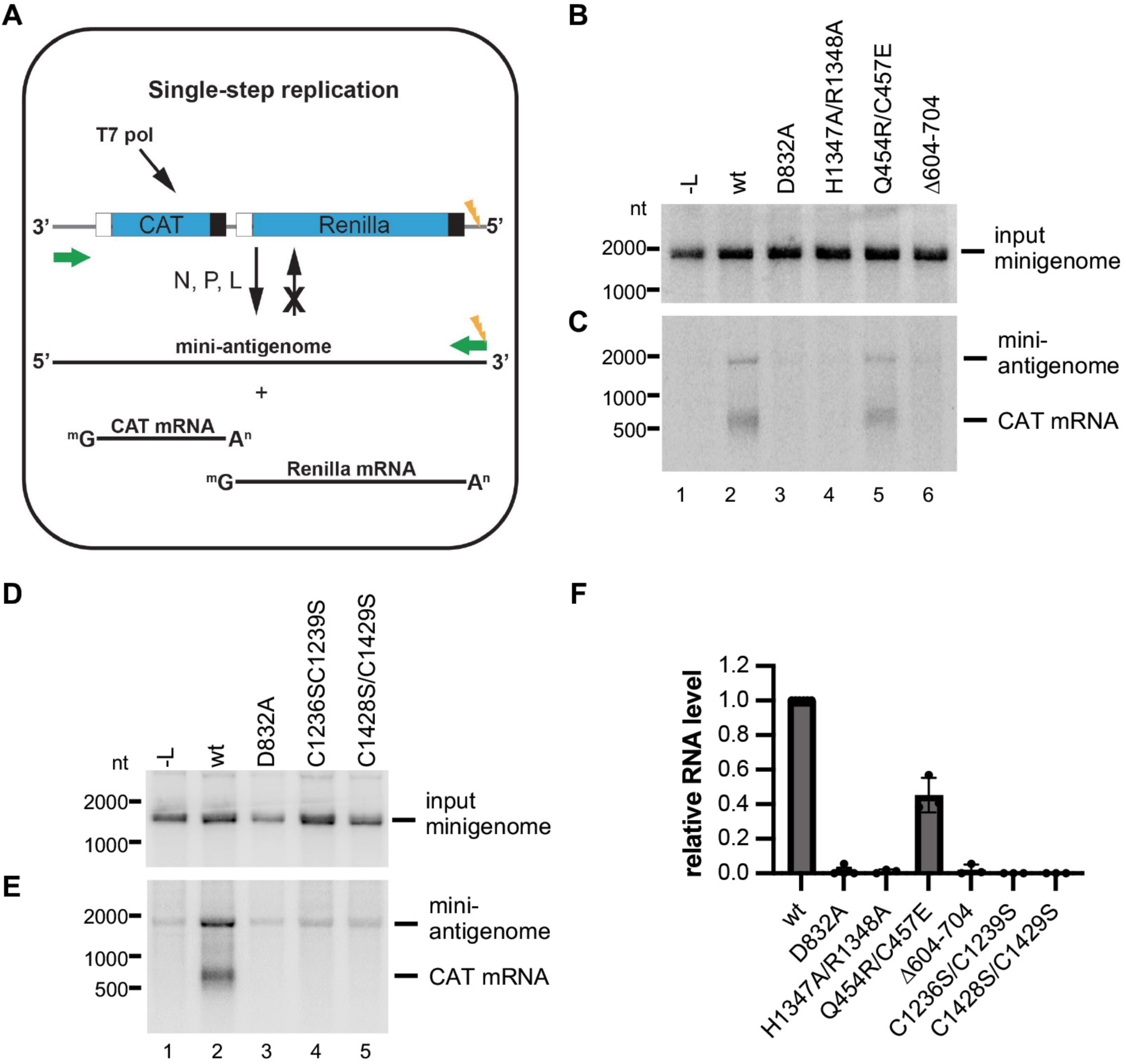
Analysis of NiV L mutants using single-step replication minigenome system to dissect effects on transcription. (A) Schematic diagram illustrating the structure of the minigenome and the products that are generated during RNA replication and mRNA transcription, as described in Figure 5D. This replication-deficient minigenome contains a mutation in the promoter region of the mini-antigenome (indicated with a lightening symbol), which restricts RNA replication to the mini-antigenome synthesis step. (B–E) Northern blot analysis of RNAs generated in the minigenome system. Panels B and D show negative sense minigenome template RNA generated by T7 RNA polymerase in each transfection reaction. Panels C and E show Northern blot analysis of positive sense mini-antigenome RNA and CAT mRNA generated by the NiV polymerase. (F) Quantification of levels of CAT mRNA from replicates of the Northern blots shown in C and E. Each bar represents the mean and standard deviation derived from three independent experiments.

## Discussion

NiV depends on its L–P polymerase complex to perform all the enzymatic activities required to generate capped and polyadenylated mRNAs and replicative RNAs. In addition, because the viral RNA template is encased within an RNP, the NiV polymerase must also be capable of transiently removing N protein from the template to feed the template RNA into its active site. And because newly synthesized replicative RNAs are encapsidated as they are synthesized, the polymerase complex is thought to capture and deliver soluble N protein onto newly synthesized replicative RNAs. The findings presented here shed light on each of these steps.

The findings presented here show that the NiV polymerase complex has similar structural features as other related polymerase structures but has some distinguishing features. The NiV L– P complex structure reveals a core comprised of the L protein RdRp and CAP domains associated with tetrameric P and one P XD. Although overall the L RdRp-CAP domain core is visualized almost in its entirety in the cryo-EM structure, suggesting that it is relatively rigid, some regions within this core could not be visualized, consistent with these features being flexible. These features include the priming and intrusion loops, which adopt different conformations in different polymerase structures, and which are less ordered in the NiV L protein structure than in other paramyxovirus L protein structures (Figure S5A–C). It is thought that the different positions adopted by these loops in different nsNSV polymerases represents different functional states. However, it is not clear what functional state is represented in most cases. In the case of the NiV polymerase structure presented here, it might represent a pre-initiation mode, in which the priming and intrusion loops have yet to be anchored into place following polymerase interactions with RNA and NTPs, or an elongation mode in which their flexibility allows nascent RNA to be extruded unimpeded. Another flexible structure is the palm insert (amino acids 600–710), which is close to the putative transcript exit channel (Figure 2C). Previously, it has been suggested that nascent mRNA and replicative RNA might emerge from different exit channels created as the globular domains adopt different conformations^33,34^. If so, this palm insert could be important for regulating the polymerase between transcription and replication. Studies with other paramyxovirus polymerases have shown this region is significant for polymerase activity but did not distinguish if it affected transcription and/or replication^37,42^. Here we add additional detail showing that deletion of the palm insert completely inhibited both mRNA and antigenome production, demonstrating that in the case of NiV, this region plays a key role during both transcription and RNA replication. Interestingly, the corresponding palm insert region of the human parainfluenza virus 3 (PIV3) L protein is also largely disordered, but the C-terminal segment of the region could be visualized as a β-strand that augments a β-sheet in the CTD (“β-latch”) and may restrict the conformation of the L C-terminal globular domains^37^. In the NiV L protein structure, the palm insert is too flexible to be detected and the L C-terminal globular domains are not visible; by analogy to the PIV3 L structure, however, the C-terminal region of the NiV L RdRp palm insert could interact with the C-terminal globular domains.

The findings presented here demonstrate the multifunctional nature of the NiV L protein CAP domain. By analogy with the rhabdovirus polymerases, the CAP domain contains residues necessary for the GTPase and PRNTase steps of cap addition^12^ and is expected to be required for mRNA transcription. However, studies with other nsNSV polymerases have shown that the HR motif, which catalyzes the PRNTase reaction also plays a role in RNA replication (which does not involve a cap addition step)^37,43^. Consistent with these findings, substitutions in the NiV L HR motif completely inhibited replication in addition to transcription. The role of the HR motif in RNA replication remains unclear, but in the RSV polymerase, substitution in this motif inhibits elongation of the replicative RNA in the promoter region^43^. Here, we also examined the significance of two zinc finger motifs, which have been observed in other nsNSV polymerases and are conserved (Figure S10), but to our knowledge, have not been functionally analyzed. As noted above, the Cys2His2 zinc finger appears to aid structural integrity of the L protein. However, mutations in both zinc finger motifs reduced the relative amounts of a truncated form of L, which is presumably generated by cleavage at the palm insert (Figure S6). This observation could suggest direct or long-range interactions between the zinc finger motifs and the palm insert, or that the zinc finger motifs influence the conformational dynamics of the RdRp core. Substitutions in each of the zinc finger motifs inhibited replication in addition to transcription, raising the possibility that they play a role common to both processes. Intriguingly, a structure of the VSV polymerase in association with RNP within virions showed that it is oriented such that the zinc finger motifs of the capping domain are proximal to the N protein of the RNP^32^. Thus, it is possible that the zinc finger motifs aid polymerase association with the RNP template.

Our findings might also provide insight into the mechanisms by which the polymerase interfaces with the RNP template. A distinction between the NiV polymerase structure reported here and other paramyxovirus polymerase structures is a visible region of P4 which snakes along the surface of L on the opposite face of the polymerase from P2 XD and closer to the template entry channel. A double amino acid substitution to disrupt the L interaction with P4 and reduced transcription and RNA replication to ∼ 25% of wild-type levels, suggesting that this L–P interface plays an equivalent role in both processes. Both transcription and RNA replication are dependent on the polymerase being able to associate with the template RNP complex, dissociate N protein from the RNA and feed the RNA into the template entry channel. Previous reports proposed an RNA synthesis mechanism in which the P2 XD domain binds to the RdRp domain, and the remaining three XD domains (P1, P3 and P4 in this case) interact with the RNP complex in turn to propel the template toward the template entrance channel^33^. However, in our structure, the peptide extension of P4 binding close to the template entrance channel (Figure 4A) suggest that P4 XD may play a particularly important role in helping to guide the RNP towards the template entrance channel.

Our findings provide new information on how the polymerase might recruit copies of free N (N^0^) for RNA encapsidation. Perhaps the most unusual feature of the complex is the ⍺-helical cap structure at the N-terminal end of the P OD. While this conformation had only been noted in X-ray crystal structures of the OD of NiV and Sendai virus^16,26,27^, it has been observed here by cryo-EM in the absence of possible crystal lattice constraints that may have influenced the conformation of this segment in prior X-ray crystal structures. Both cryo-EM 3D variability analysis and molecular dynamics analysis suggest that the tip of long coiled-coil that is capped by a bunding of helices is highly flexible. This flexibility is probably important for how P interacts with L during RNA replication, which is a dynamic process in which P negotiates interactions with N^0^ and RNP. Interestingly, P XD, which docks on the L RdRp near the nascent RNA exit channel, is also known to bind the C-terminal region of N^15^ (Figure S11). Thus, it is tempting to speculate that N^0^ copies could be recruited to the L RdRp core first through binding of P-NTD to N-terminal region of N^0^ followed by engagement of the N^0^ C-terminal region with the P2 XD binding site, thus allowing for efficient nascent RNA encapsidation during replication (Figure S11).

In summary, we have determined the structure of NiV L–P complex, characterized its properties through biophysical means, and performed functional analyses to identify features that play key roles in transcription and replication. These findings enhance our understanding of nsNSV polymerases and the mechanisms by which they engage in transcription or RNA replication and have the potential to allow for rational drug design against NiV infection.

**Figure S1.**
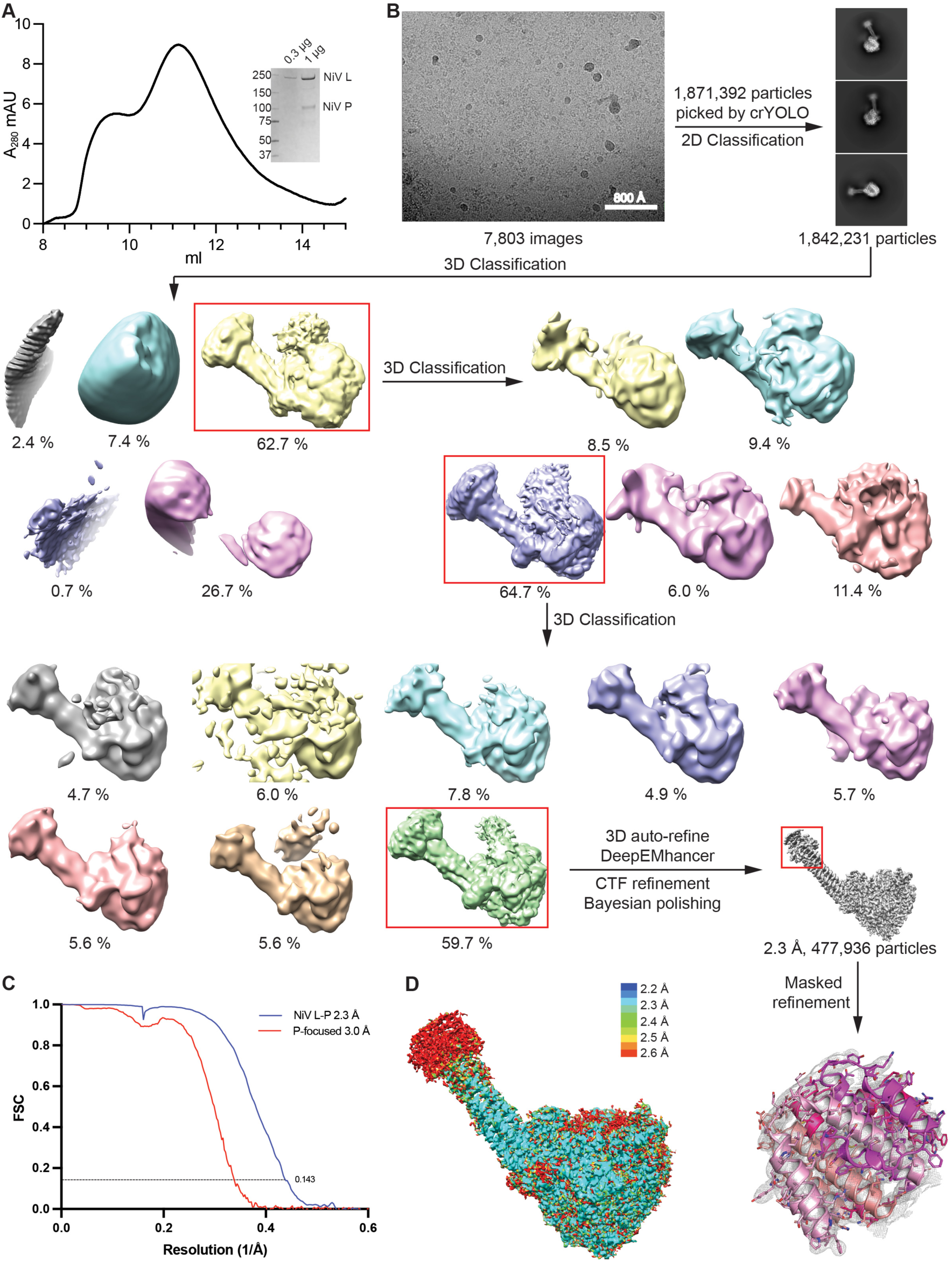
Cryo-EM reconstruction of NiV L–P complex. (A) SEC profile of NiV L–P complex using the column Superose 6^®^ Increase 10/300 GL. The complex has a retention volume around 11 ml. The purity was examined using SDS-PAGE, with a Coomassie-stained gel shown (inset). NiV L protein has a size around 260 kDa; NiV P protein has a size around 80 kDa but like other P proteins, has an apparent size that is larger than its molecular weight, around 100 kDa. (B) Workflow used for cryo-EM data processing of NiV L–P complex. (C) Fourier shell correlation (FSC) curves. The threshold used to estimate the resolution is 0.143. (D) ResMap^47^ showing local resolution of the NiV L–P complex.

**Figure S2.**
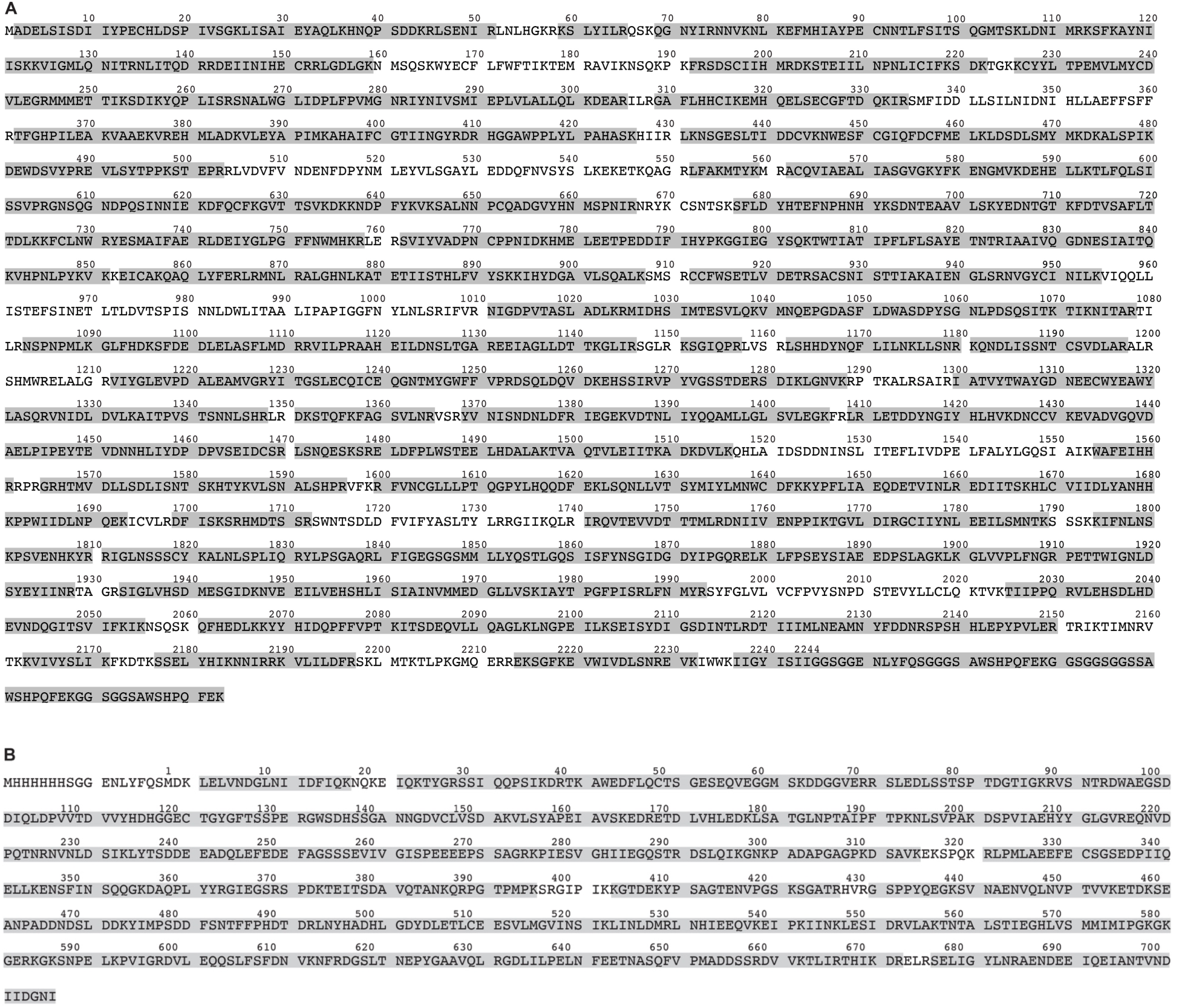
LC-MS/MS sequencing of purified NiV L and P. (A) The NiV L band that is near 250 kDa marker band on SDS-PAGE (Figure S1A) was cut and sent for trypsin digestion, followed by LC-MS/MS sequencing. Fragments of NiV L detected by LC-MS/MS are highlighted in gray. NiV L has a coverage more than 82%. The sequence C-terminal of residue 2244 is a glycine-serine linker and triple-strep tag. (B) The NiV P band near 100 kDa marker band on SDS-PAGE (Figure S1A) was cut and sent for LC-MS/MS sequencing. Fragments detected are highlighted in gray. NiV P has a coverage over 94%. The sequence N-terminal of residue 1 is a histidine tag and serine-glycine linker.

**Figure S3.**
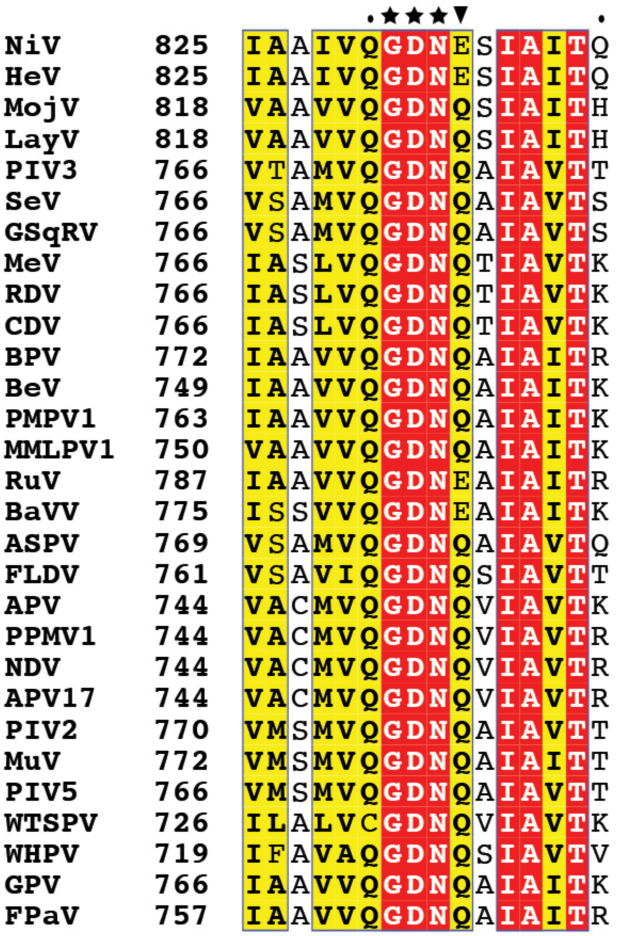
Amino acid sequence alignment of paramyxovirus L proteins focusing on the GDNQ/E motif. Residues GDN (marked by stars) are the conserved residues among all the L proteins aligned. The fourth residue, marked with a black triangle, while in most of the cases it is glutamine, is glutamate in a few sequences, including NiV L. NiV, Nipah virus; HeV, Hendra virus; MojV, Mojiang virus; LayV, Langya virus; PIV3, human parainfluenza virus 3; SeV, Sendai virus; GSqRV, giant squirrel respirovirus; MeV, measles virus; RDV, rinderpest virus; CDV canine distemper virus; BPV, bat paramyxovirus; BeV, Belerina virus; PMPV1, Pohorje myodes paramyxovirus 1; MMLPV1, mountain Mabu Lophuromys paramyxovirus 1; RuV, Ruloma virus; BaVV, bank vole virus 1; ASPV, Atlantic salmon paramyxovirus; FLDV, Fer-de-Lance paramyxovirus; APV, avian paramyxovirus UP0216; PPMV1, pigeon paramyxovirus 1; NDV, Newcastle disease virus; APV17, avian paramyxovirus 17; PIV2, human parainfluenza 2 virus; MuV, mumps virus; PIV5, parainfluenza virus 5; WTSPV, Wenling tonguesole paramyxovirus; WHPV, Wenling tonguesole paramyxovirus; GPV, gerbil paramyxovirus; FPaV, feline paramyxovirus 163. See Supplementary Table 3 for GenBank accession numbers.

**Figure S4.**
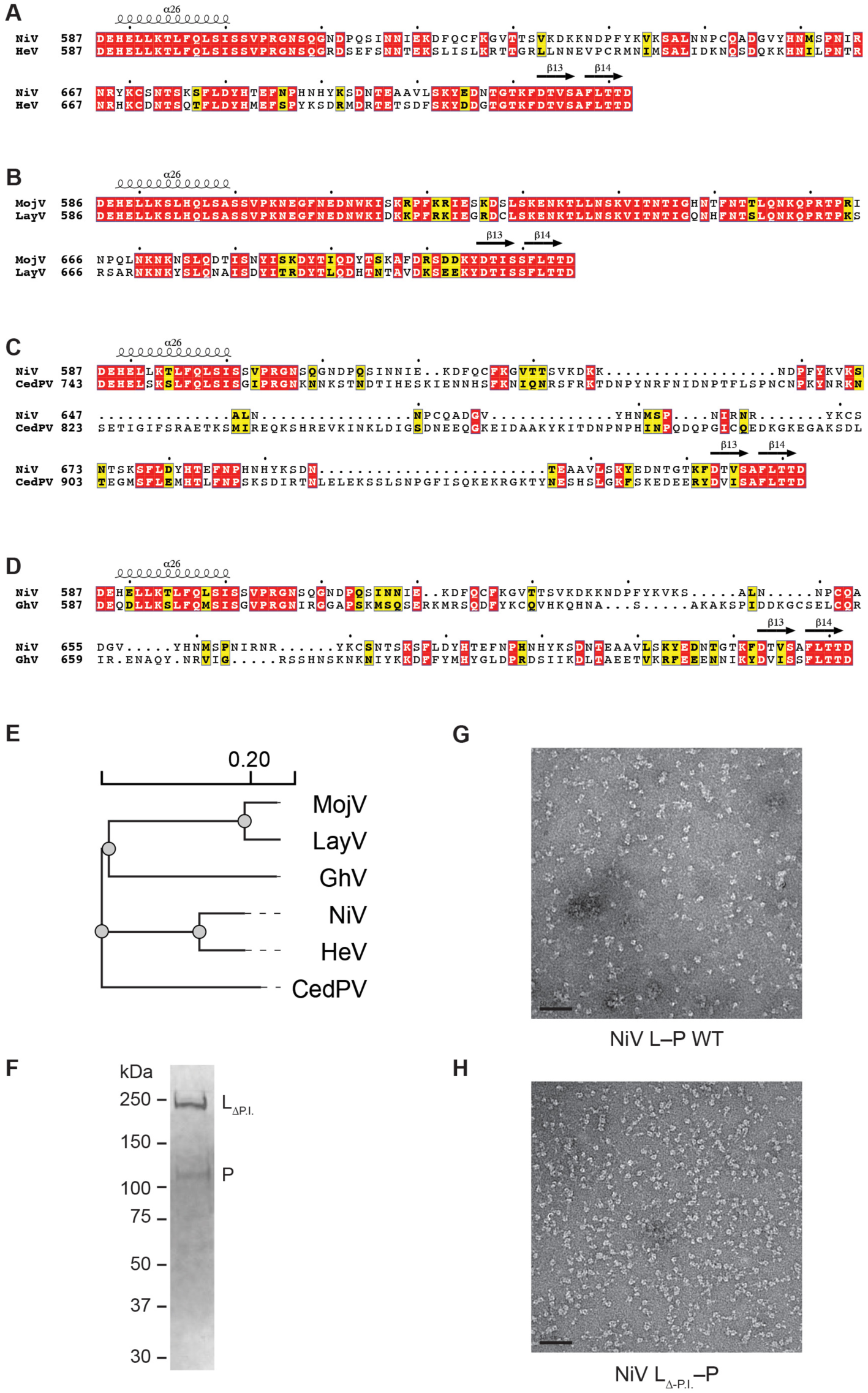
Sequence comparison of the palm insert in henipaviruses and negative stain images of recombinant NiV L–P proteins. (A) Amino acid sequence alignment of palm insert region of NiV and HeV. (B) Amino acid sequence alignment of palm insert region of MojV (Mojiang virus) and LayV (Langya virus). (C) Amino acid sequence alignment of palm insert region of NiV and CedPV (Cedar virus). Cedar virus has a particularly long palm insert. (D) Amino acid sequence alignment of palm insert region of NiV and GhV (Ghana virus). (E) Phylogenetic tree of the indicated nsNSVs based on L amino acid sequences generated using Clustal Omega^45^. Genera and families are indicated. See Table S3 for additional information. (F) SDS-PAGE gel of eluted fraction from Streptactin purification of NiV L_Δ-P.I._–P. Imaging was performed with a stain free gel system. (G) Negative stain image of NiV L–P wild-type (WT). After Streptactin purification, NiV L–P was diluted to 0.03 mg/ml for negative stain. (H) Negative stain image of NiV L_Δ-P.I._–P. After Streptactin purification, NiV L_Δ-P.I._–P was diluted to 0.06 mg/ml for negative stain. The scale bar represents 50 nm.

**Figure S5.**
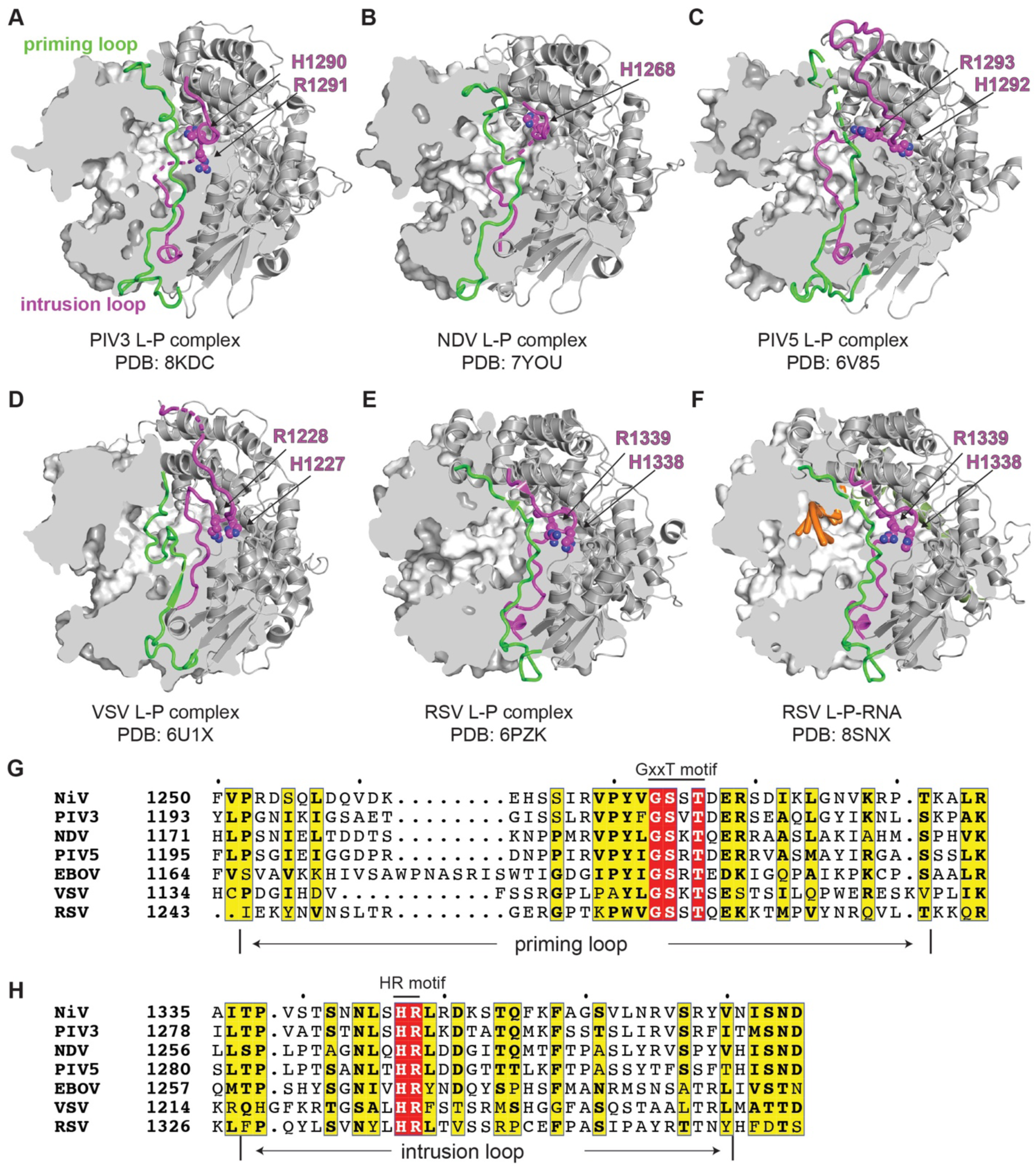
CAP domain priming loop and intrusion loop in additional polymerase structures. (A–F) Cryo-EM structures of PIV3 L–P (PDB: 8KDC) (A), NDV L–P (PDB: 7YOU) (B), PIV-5 L–P (PDB: 6V85) (C), VSV L–P (PDB: 6U1X) (D), RSV L–P (6PZK) (E) and RSV L–P-RNA (PDB: 8SNX) (F), whose L protein RdRp domains are shown in surface representation with partially clipped surfaces, and the CAP domains are shown as ribbon diagrams. The priming (green) and intrusion (purple) loops are disordered in the cryo-EM structures. The disordered portions are shown as dashed lines. The side chains of intrusion loop HR residues are shown as spheres and indicated. Nucleic acids are shown in orange. Except for NDV, whose R1269 was in a disordered segment, the HR residues of these structures are shown. (G–H) Amino acid sequence alignment of NiV, PIV-3, NDV, PIV-5, EBOV, VSV, and RSV, focusing on the priming loop (G) and intrusion loop (H) regions.

**Figure S6.**
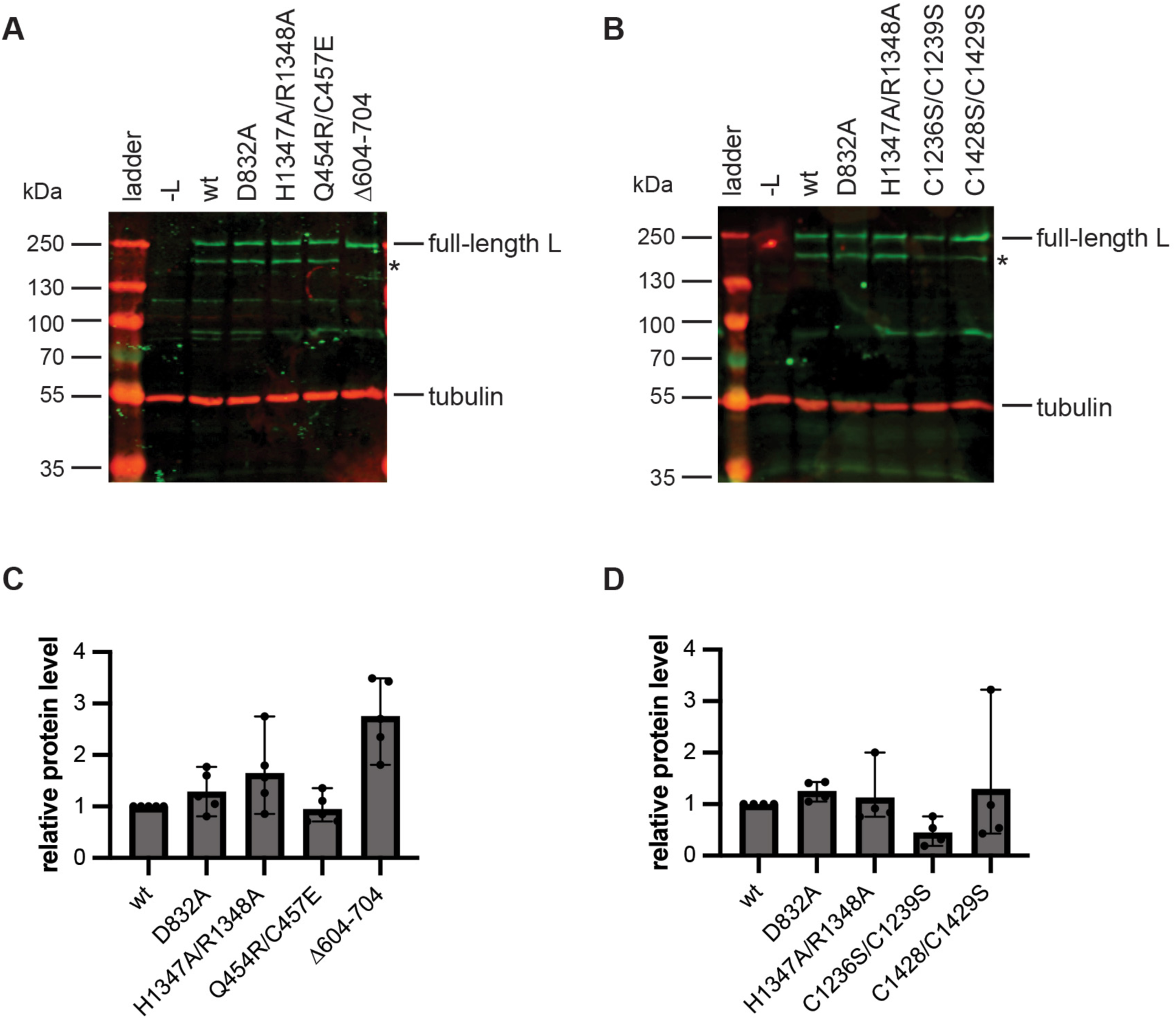
Confirmation of mutant L protein expression in the minigenome system. (A–B) Western blot analysis of minigenome transfected cell lysates probed with a strep-tag specific antibody to detect L; tubulin was detected with a specific antibody as a loading control. The band marked with an asterisk is the appropriate molecular weight to be truncated L protein that had been cleaved in the palm insert. (C–D) Quantification of the levels of non-truncated L protein from replicates of the Western blots shown in A and B. Each bar represents the mean and standard deviation derived from three independent experiments.

**Figure S7.**
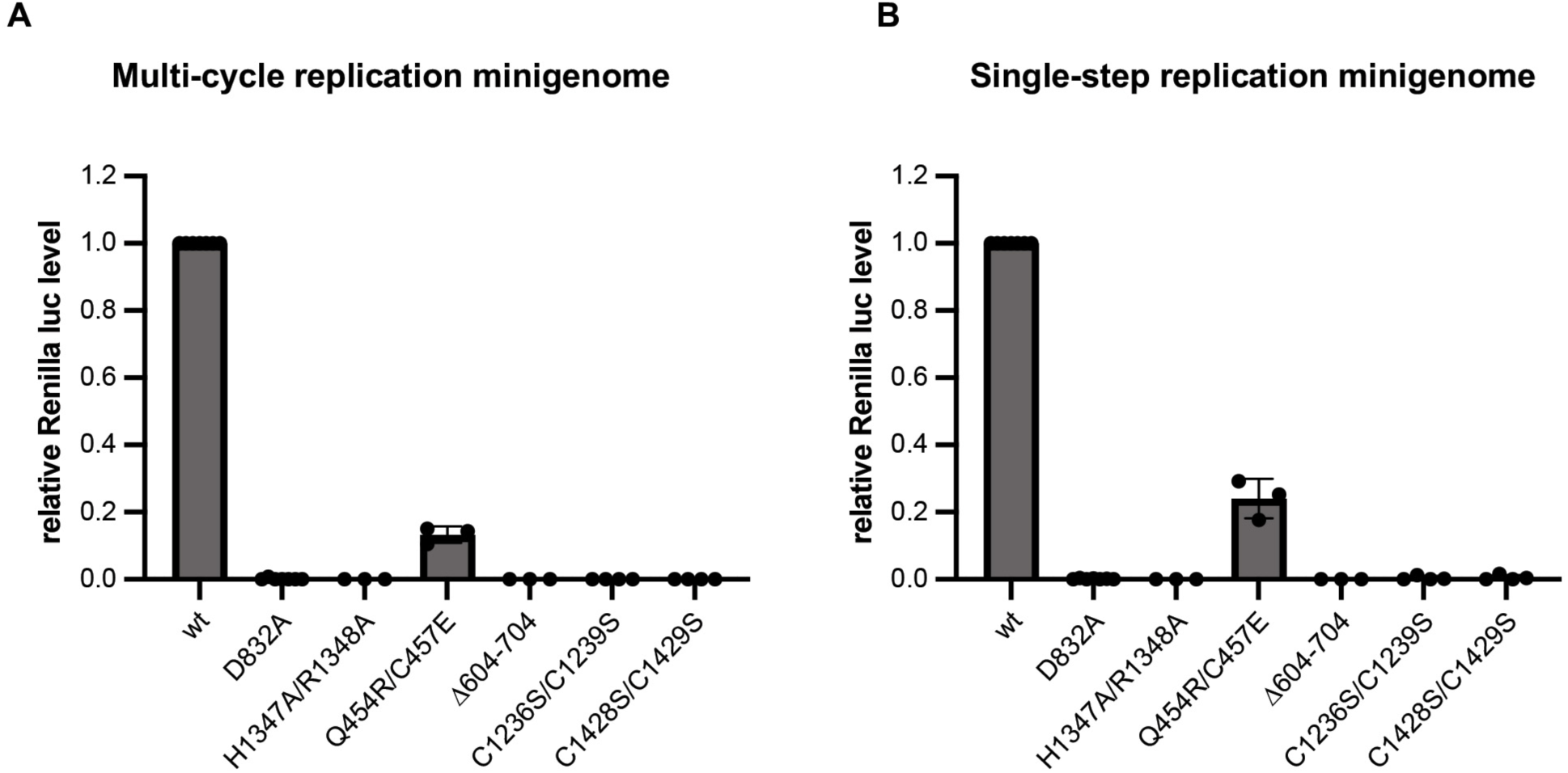
RNA synthesis activities of mutant L proteins measured by luciferase activity. (A–B) Renilla luciferase activity of L mutants in an assay with a multi-cycle replication minigenome (A) or single-step replication minigenome (B). In addition to cells being transfected with plasmids expressing minigenome RNA, and N, P and wild-type (wt) or mutant L proteins, cells were co-transfected with a T7 driven plasmid expressing firefly luciferase as a control for transfection efficiency. Renilla luciferase values were normalized to firefly luciferase values, the background signal present in the D832A sample was subtracted from each value, and the values were normalized to L wt. The bars show the mean and SD from three independent experiments.

**Figure S8.**
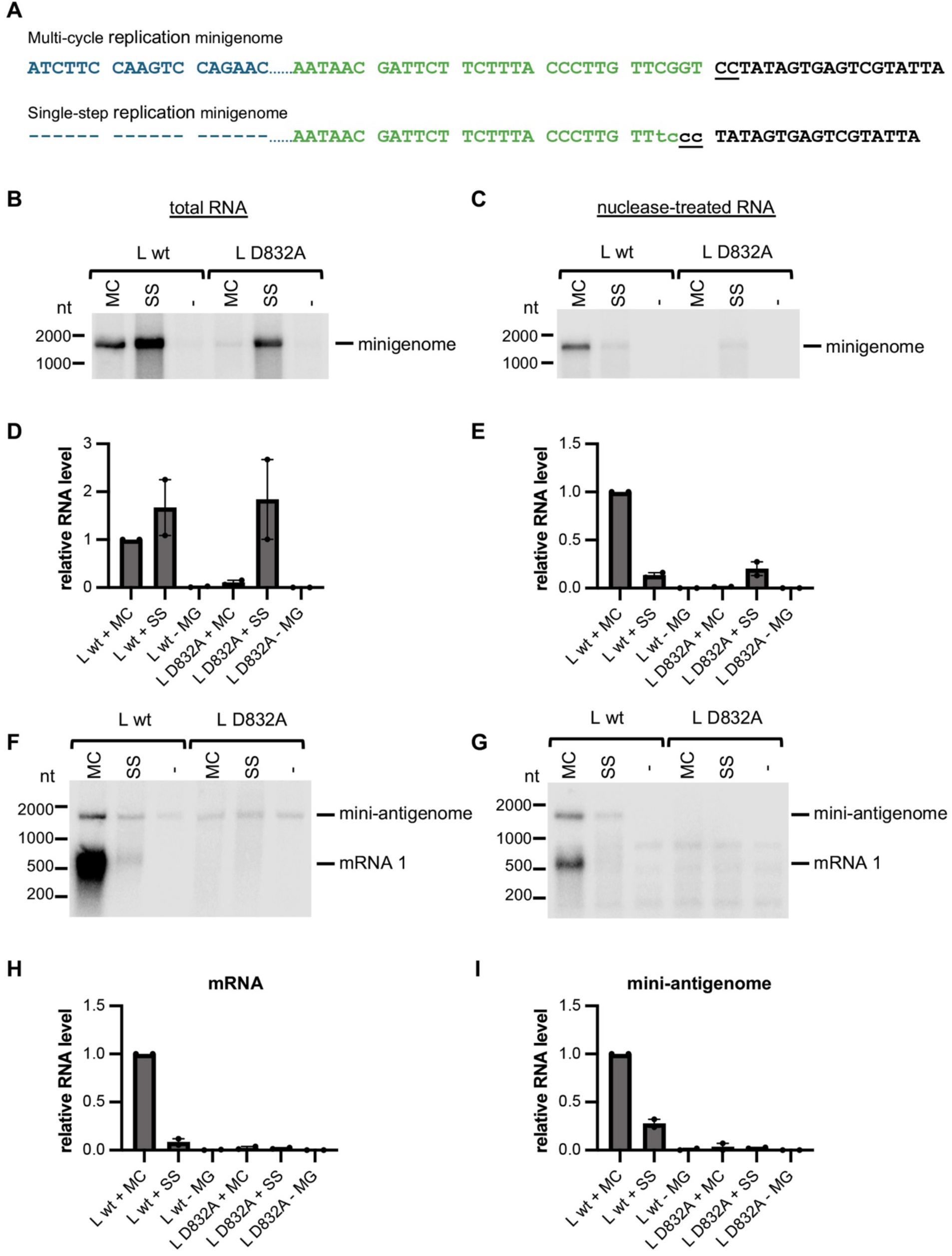
Validation of a single-step replication minigenome assay. (A) Sequence alignment showing the changes introduced to increase T7 promoter efficiency and ablate the promoter at the 3’ end of the NiV antigenome. The sequence is shown as positive (antigenome) sense RNA. The internal NiV promoter element (promoter element 2), which was deleted to limit the minigenome to a single-step of replication is shown in blue, the complement of the trailer region is shown in green, and the T7 promoter is shown in black. The T7 RNA polymerase initiates opposite the C residues that are underlined to synthesize negative sense RNA. Substitutions introduced into the trailer promoter are shown in lowercase. The multi-cycle replication minigenome followed the “rule of six” (i.e., its nucleotide length was divisible by six) except for the additional G residues contributed by the T7 promoter; the single-step replication minigenome followed the “rule of six” including the additional G residues contributed by the T7 promoter. (B–C) Northern blot analysis of negative sense (minigenome sense) RNA generated by either T7 RNA polymerase alone, or T7 RNA polymerase and NiV polymerase. The blots show RNA generated in cells transfected with plasmids expressing either wt L or L with a substitution in the GDNE motif of the RdRp (L D832A). Cells were transfected with plasmids expressing either multi-cycle (MC) or single-step (SS) replication minigenome, or minigenome (MG) was omitted from the transfection (-). Panel B shows total RNA harvested from transfected cells. Panel C shows RNA from a parallel transfection in which the cell lysate was treated with micrococcal nuclease prior to RNA purification to reduce levels of unencapsidated RNA (and distinguish encapsidated minigenome template RNA). (D–E) Quantification of the bands shown in B and C, respectively. (F–G) RNA from the same transfections as used for B and C, in which the Northern blot was probed to detect positive sense RNA (mini-antigenome and CAT mRNA). (H–I) Quantification of the bands shown in F and G, respectively. Panel H shows quantification of CAT mRNA, panel I shows quantification of nuclease-resistant antigenome RNA. The data shown in panels B, C, F and G are representative of two independent experiments. The bars in panels D, E, H and I show the two data points and the mean for the two independent experiments, with normalization to L wt + MC minigenome in each case. This experiment shows that the single-step replication minigenome had a stronger T7 promoter than the multi-cycle replication minigenome (panel B and D). However, the single-step replication minigenome was not amplified by wt L protein (panel C and E) and consequently produced relatively low levels of CAT mRNA (panels F and H) and mini-antigenome (panels G and I).

**Figure S9.**
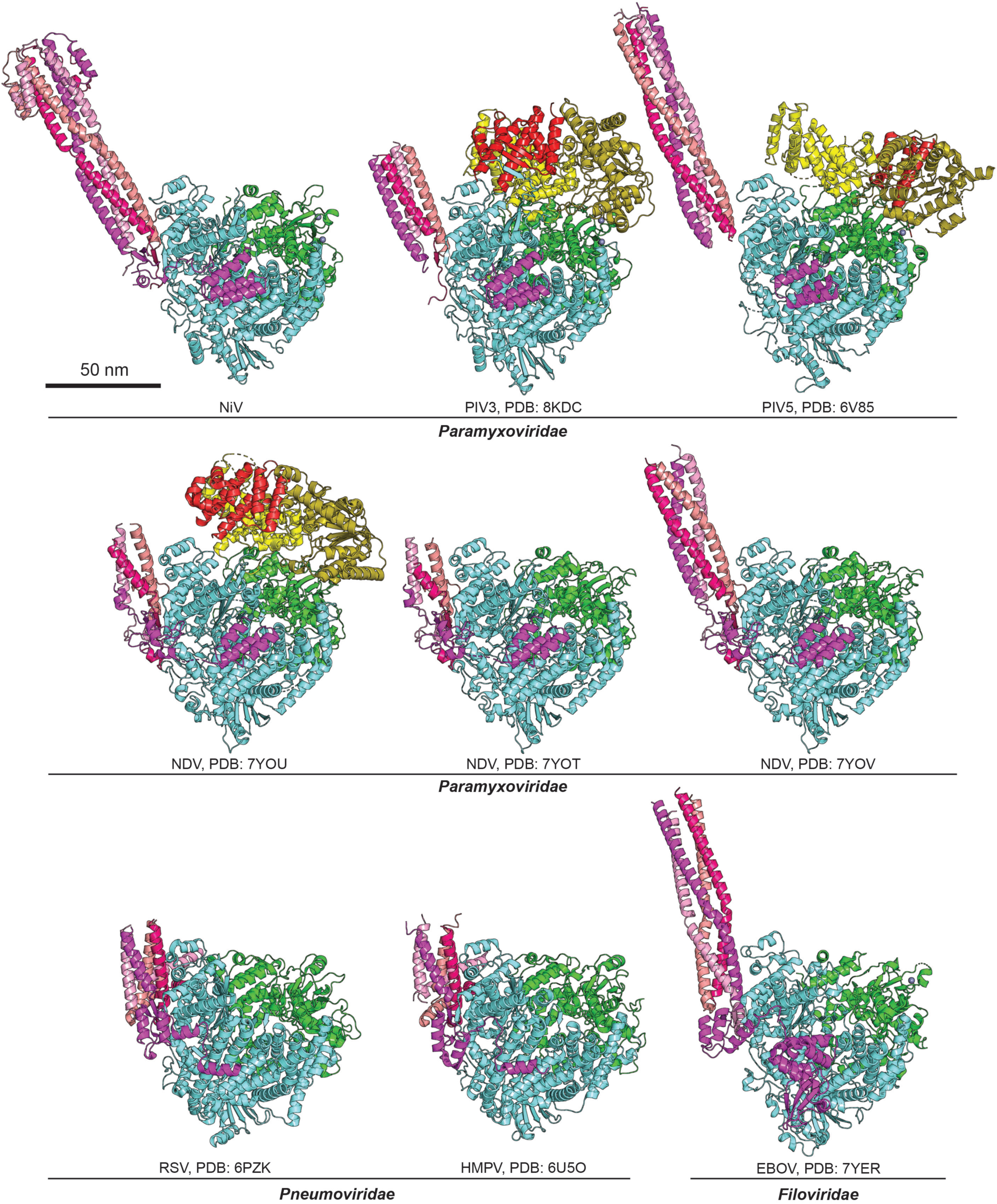
Comparison of nsNSV cryo-EM structures. Cryo-EM structures of NiV, PIV-3 (PDB: 8KDC), PIV-5 (PDB: 6V85), and NDV (PDB: 7YOV, 7YOT and 7YOV) in the family *Paramyxoviridae*, RSV (PDB: 6PZK) and HMPV (PDB: 6U5O) in the family *Pneumoviridae*, and EBOV (PDB: 7YER) in the family *Filoviridae* show part of the P proteins. NDV has three structures from the same dataset showing different lengths of P OD domain. Structures of PIV-3, PIV-5, and NDV include the CD-MTase-CTD domains (colored light yellow, dark yellow, and red, respectively), but PIV-5 has a different arrangement of these domains from PIV-3 and NDV. Only NiV P has an ⍺-helical cap structure in the P N-terminal region of the P OD. Metals are shown as gray spheres.

**Figure S10.**
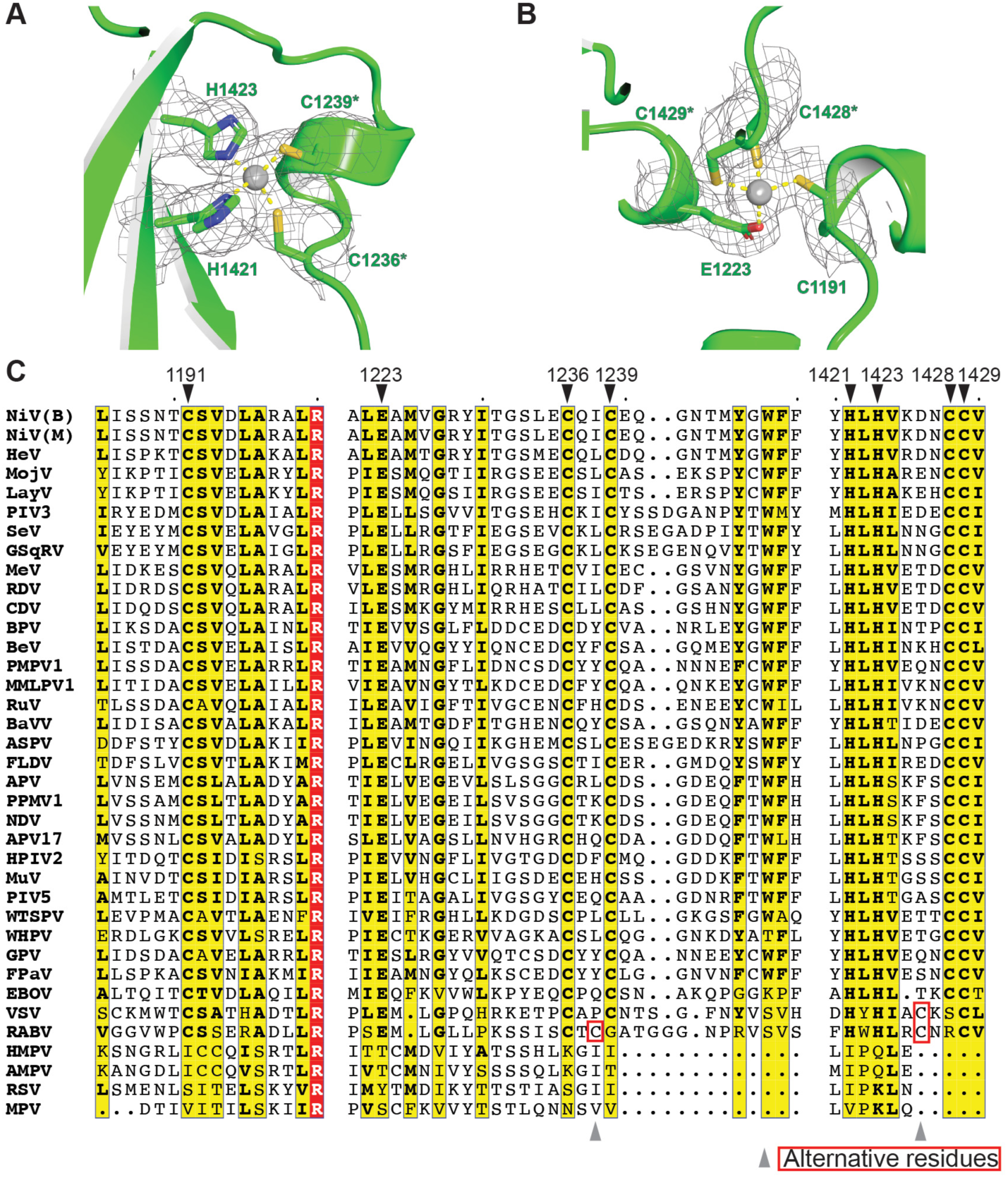
CAP domain zinc fingers. (A–B) Two zinc fingers are observed in NiV L–P complex, one in Cys2His2 fold group (A) and the other in Gag Knuckle fold group (B). cryo-EM density for the zinc metal (gray sphere) and neighboring side chains are shown. (C) Amino acid sequence alignment of nsNSVs showing the residues forming zinc fingers. While most of these sequences have conserved residues (indicated by inverted black triangles, the numbering is based on NiV L residues), pneumoviruses including HMPV, AMPV, RSV and MPV do not have all of these conserved residues and rhabdoviruses like VSV and RABV may have alternative residues (highlighted in red boxes) that are near the conserved residues (indicated by gray triangles below the sequence alignment). NiV (B), Nipah virus, Bangladesh strain; NiV (M), Nipah virus, Malaysian strain; HeV, Hendra virus; MojV, Mojiang virus; LayV, Langya virus; GhV, Ghana virus; PIV3, human parainfluenza 3; SeV, Sendai virus; GSqRV, giant squirrel respirovirus; MeV, measles virus; RDV, rinderpest virus; CDV canine distemper virus; BPV, bat paramyxovirus; BeV, Belerina virus; PMPV1, Pohorje myodes paramyxovirus 1; MMLPV1, mountain Mabu Lophuromys paramyxovirus 1; RuV, Ruloma virus; BaVV, bank vole virus 1; TPMV, Tupaia paramyxovirus; ASPV, Atlantic salmon paramyxovirus; FLDV, Fer-de-Lance paramyxovirus; APV, avian paramyxovirus UP0216; PPMV1, pigeon paramyxovirus 1; NDV, Newcastle disease virus; APV17, pigeon paramyxovirus 17; HPIV2, human parainfluenza 2 virus; MuV, mumps virus; PIV5, parainfluenza virus; WTSPV, Wenling tonguesole paramyxovirus; WHPV, Wenling tonguesole paramyxovirus; GPV, gerbil paramyxovirus; FPaV, feline paramyxovirus 163; EBOV, Ebola virus; VSV, vesicular stomatitis virus; RABV, rabies virus; HMPV, human metapneumovirus; AMPV, avianmetapenumovirus; RSV, respiratory syncytial virus; MPV, murine pneumonia virus.

**Figure S11.**
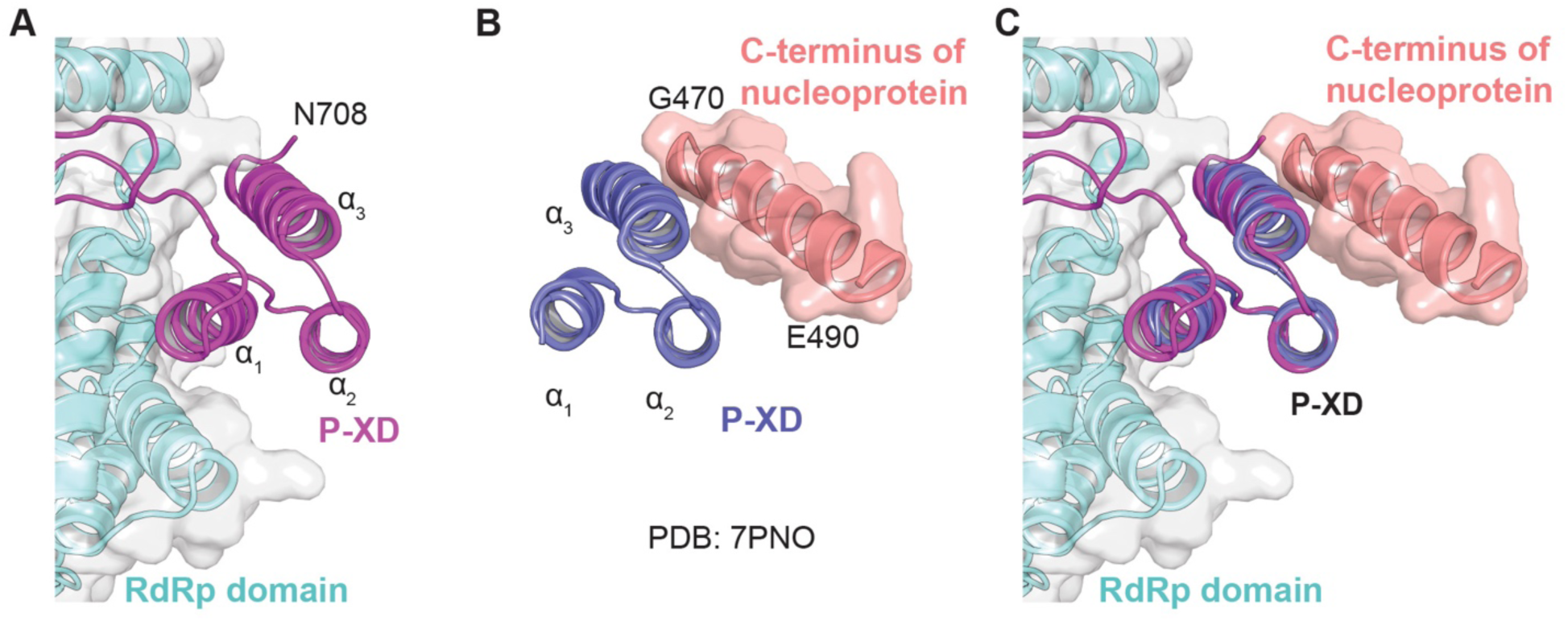
Comparison of NiV P XD in complex with NiV L RdRp domain and the C-terminus of nucleoprotein. (A) NiV P XD in complex with NiV L RdRp domain. P XD folds into three *α*-helices, *α*-1–3, among which, *α*-1 and *α*-3 interact with the NiV L RdRp domain. The last residue, N708, of NiV P XD domain in the structure is indicated. (B) Crystal structure of NiV P XD in complex with a C-terminal fragment of NiV N (residues G470–E490 are modeled) (PDB: 7PNO)^15^. P XD also folds into three *α*-helices but *α*-2 and *α*-3 interact with the C-terminal fragment of NiV N. (C) A model showing a possible interaction of NiV L RdRp domain, P XD, and NiV N, based on structural superposition of the NiV L–P cryo-EM structure and the previously reported crystal structure of NiV P XD in complex with the C-terminal fragment of NiV N (PDB: 7PNO)^15^.

**Table S1.** Data collection, image processing, and model refinement.

|  |  |
| --- | --- |
|  | NiV L–P |
| **Data collection and processing** |  |
| Microscope | Titan Krios |
| Detector | K3 Summit Direct Electron Detector |
| Magnification | 105,000 x |
| Voltage (kV) | 300 |
| Electron exposure (e<sup>-</sup>/Å<sup>2</sup>) | 53 |
| Defocus range (μm) | -1.0 to -2.5 |
| Pixel size (Å) | 0.83 |
| Symmetry imposed | C1 |
| Initial particles | 1,871,392 |
| Final particle images (no.) | 477,936 |
| Map resolution (Å) | 2.3 |
| FSC threshold | 0.143 |
| **Model refinement and validation** |  |
| Initial model used | L: Phyre2 prediction; P: 4N5B |
| R.m.s.deviation |  |
| Bond lengths (Å) | 0.006 |
| Bond angles (°) | 0.81 |
| Validation |  |
| Clashscore | 13 |
| Favored (%) | 96.3 |
| Allowed (%) | 3.6 |
| Outliers (%) | 0.1 |

**Table S2.** Alignment of NiV L with other NNS virus L proteins.

| **Virus** | **PDB** | **residues** | **RMSD of Cα (# of aligned atoms)** |
| --- | --- | --- | --- |
| HMPV | 6U5O | 8–1380 | 3.97 (777) |
| RSV | 6PZK | 10–1460 | 4.20 (819) |
| VSV | 5A22 | 35–1334 | 4.59 (825) |
|  | 6U1X | 35–1332 | 4.66 (826) |
| RABV | 6UEB | 29–1352 | 4.39 (741) |
| EBOV | 7YER | 8–1383 | 3.15 (778) |
|  | 7YES | 8–1383 | 3.37 (794) |
| PIV5 | 6V85 | 5–1397 | 1.89 (907) |
|  | 6V86 | 5–1397 | 1.91 (905) |
| NDV | 7YOU | 8–1384 | 1.54 (985) |
|  | 7YOV | 8–1384 | 1.54 (981) |
|  | 7YOT | 8–1384 | 1.49 (973) |
| PIV3 | 8KDB | 8–1395 | 1.84 (1023) |
|  | 8KDC | 10–1395 | 1.86 (1031) |

**Table S3.** GenBank accession number for L sequences.

| **Virus** | **Name** | **GenBank accession number** |
| --- | --- | --- |
| NiV (B) | Nipah virus, Bangladesh strain | AAY43917.1 |
| NiV (M) | Nipah virus, Malaysia strain | NP_112028.1 |
| HeV | Hendra virus | NP_047113.3 |
| CedPV | Cedar virus | YP_009094087.1 |
| GhV | Ghana virus | YP_009091839.1 |
| MojV | Mojiang virus | YP_009094096.1 |
| LayV | Langya virus | UUV47207.1 |
| PIV3 | Human parainfluenza virus 3 | WCF97250.1 |
| SeV | Sendai virus | YP_010790275.1 |
| GSqRV | Giant squirrel respirovirus | YP_010806263.1 |
| MeV | Measles virus | NP_056924.1 |
| RDV | Rinderpest virus | P41357.1 (UniProtKB) |
| CDV | Canine distemper virus | NP_047207.1 |
| BPV | Bat paramyxovirus | YP_010796325.1 |
| BeV | Belerina virus | YP_010801294.1 |
| PMPV1 | Pohorje myodes paramyxovirus 1 | YP_010085024.1 |
| MMLPV1 | Mountain Mabu Lophuromys virus 1 | YP_009666847.1 |
| RuV | Ruloma virus | YP_010801062.1 |
| BaVV1 | Bank vole virus 1 | YP_010085015.1 |
| TPMV | Tupaia paramyxovirus | NP_054697.1 |
| ASPV | Atlantic salmon paramyxovirus | YP_009094152.1 |
| FLDV | Fer-de-Lance paramyxovirus | NP_899661.1 |
| APV | Avian paramyxovirus UP0216 | YP_009508504.1 |
| PPMV1 | Pigeon paramyxovirus 1 | QBZ96492.1 |
| NDV | Newcastle disease virus | YP_009513199.1 |
| APV17 | Avian paramyxovirus 17 | YP_010796382.1 |
| HPIV2 | Human parainfluenza virus 2 | NP_598406.1 |
| MuV | Mumps virus | NP_054714.1 |
| PIV5 | Parainfluenza virus 5 | QWL54719.1 |
| WTSPV | Wenling tonguesole paramyxovirus | YP_010790495.1 |
| WHPV | Wenling hoplichthys paramyxovirus | YP_010790504.1 |
| GPV | Gerbil paramyxovirus | UWK09075.1 |
| FPaV | Feline paramyxovirus 163 | YP_010799262.1 |
| EBOV | Zaire ebola virus | AQA27307.1 |
| VSV | Vesicular stomatitis virus | NP_041716.1 |
| RABV | Rabies virus SAD-B19 | P16289.1 |
| HMPV | Human metapneumovirus | YP_009513273.1 |
| AMPV | Avian metapneumovirus | Q2Y2L8.2 (UniprotKB) |
| RSV | Respiratory syncytial virus | NP_044598.1 |
| MPV | Murine pneumonia virus | YP_010774683.1 |

## Methods

### Cloning and insect cell expression of L–P complexes

The coding sequence of NiV L (GenBank: AAY43917.1) and P (GenBank: AAY43912.1) were synthesized and codon-optimized for Bac-to-Bac expression system using pFastBac Dual transfer vector. The sequences of NiV L and P were fused with a C-terminal triple-strep tag after a TEV protease cleavage site and an N-terminal His_6_ tag followed by a TEV protease cleavage site, respectively. NiV L was inserted downstream of the polyhedrin promoter and P was inserted downstream of the p10 promoter in the pFastBac Dual. Recombinant bacmid containing NiV L and P genes was isolated after transposing into DH10Bac competent cells following the user guide of the Bac-to-Bac Baculovirus Expression System (Invitrogen). Viral stock generated from purified bacmids was amplified and used for protein expression. Two liters of *Spodoptera frugiperda* 9 (Sf9) at a density of 2.5×10^6^ cells/ml were infected with amplified viruses to co-express NiV L and P proteins at 27°C for 72 h.

### NiV L–P complex protein purification

The pellet of Sf9 cells expressing the NiV L–P complex was resuspended in the lysis buffer (50 mM Tris, pH 7.5, 500 mM NaCl, 10% (v/v) glycerol, 6 mM MgSO_4_, 1 mM dithiothreitol (DTT), 1% (v/v) Triton X-100) supplemented with cOmplete^TM^, EDTA-free protease inhibitor cocktail (Roche). After high-speed centrifugation at 20,000 rpm for 2 h at 4 °C, the supernatant containing the target proteins was incubated with Strep-Tactin^®^ XT Sepharose resin (IBA Lifesciences) in cold room (4°C) for over 1 h and then loaded onto a gravity column. The beads were washed using buffer A (50 mM Tris pH 8.0, 500 mM NaCl, 10% glycerol (v/v), 2 mM MgSO_4_, 1 mM DTT), and the bound proteins were eluted using 50 mM biotin in buffer A. The eluates were analyzed by sodium dodecyl sulfate-polyacrylamide gel electrophoresis (SDS-PAGE) and the putative L and P bands were subjected to LC-MS/MS analysis (Harvard Center for Mass Spectrometry) to confirm their identities. The fractions containing L–P protein were further purified using a size-exclusion column (Superose 6 Increase 10/300 GL, GE healthcare) equilibrated with size exclusion chromatography (SEC) buffer (20 mM Tris, pH 7.5, 500 mM NaCl, 2 mM MgSO_4_, 1 mM DTT). The peak fractions near 11 mL were pooled and concentrated. The protein homogeneity was characterized by negative-stain EM as well as mass photometry (Refeyn Two^MP^) (see below).

### Mass Photometry analysis of NiV L–P protein

Mass photometry analyses were carried out with a Refeyn Two^MP^ mass photometer (Refeyn LTD, Oxford, UK) at room temperature. Glass coverslips and gaskets were cleaned with HPLC-grade water and isopropanol and dried under filtered gas before use. NiV L–P was diluted to 200 nM with the SEC buffer. Eighteen microliters of buffer were used to find the camera focus prior to loading 2 µl of the sample onto the gasket. Acquisition camera image size was set to “medium”. Data is collected as a 1 min movie and then processed using ratiometric imaging. To correlate ratiometric contrast to mass, the Refeyn Two^MP^ instrument was calibrated using molecular standards of BSA monomer (66 kDa), BSA dimer (132 kDa) and thyroglobulin (MW 660 kDa) with a molecular weight error less than 5%. The experiment was performed twice. Data were analyzed using Discover^MP^ software (Refeyn LTD, Oxford, UK).

### In vitro RNA synthesis assay

L protein was quantified by SDS-PAGE and densitometric comparison with a BSA standard curve on the same gel. L–P complexes (8–90 nM final concentration) were incubated with 2 μM NiV *le* 1–17 template with the sequence 3’-UGGUUUGUUCCCUUUUA-5’ in a buffer containing 50 mM Tris-HCl, pH 7.5, 8 mM MgCl_2_, 5 mM DTT and 10% glycerol (v/v) for 10 min at 30°C. Reactions were initiated by the addition of ATP and CTP to a final concentration of 1 mM each and 10 μCi of [α-^32^P]-GTP tracer (Revvity, 3000 Ci/mmol and 10 μCi/μl, final concentration 100–150 nM) in a total reaction volume of 50 μl and allowed to proceed for 1 h at 30°C. The polymerase complex was inactivated by heating to 95°C for 5 min. Reactions were subsequently treated with 10 U of calf intestinal alkaline phosphatase (NEB, 10,000 U/ml) for 1 h at 37°C. 200 μl of 0.25% SDS in nuclease-free water were added to each and RNA products were isolated by extraction with 250 μl of acid-phenol:chloroform (Invitrogen). After the addition of 170 mM NaCl and 15 μg glycogen (Invitrogen), RNA products were precipitated with ethanol overnight at −20°C. Pellets were washed once with ice-cold 70% (v/v) ethanol, briefly air-dried and resuspended in 8 μl water. An equal volume of 2 x STOP buffer (20 mM EDTA, 0.01% each bromophenol blue and xylene cyanol in deionized formamide) was added before heat denaturing samples for 5 min at 95°C. Samples were flash cooled on ice and loaded onto a 20% acrylamide sequencing gel containing 7 M urea in Tris-borate-EDTA buffer. Gels were vacuum dried at 80°C onto Whatman 3MM paper. RNA products were visualized by phosphor imaging (Typhoon IP, GE Healthcare Life Technologies).

### Cryo-EM sample preparation and data collection

An aliquot of 4 µl of NiV L–P complex at 0.7 mg/ml was applied to a freshly glow-discharged Quantifoil Cu 1.2/1.3 400 mesh grid. The sample was blotted for 3 s after incubation for 15 s at 4°C with a relative humidity of 100%, then plunge-frozen in liquid ethane using Vitrobot Mark IV (Thermo Fisher Scientific, USA), and stored in liquid nitrogen. Cryo-EM data were collected on a Titan Krios microscope (Thermo Fisher Scientific, USA) under 300 kV, equipped with a K3 Summit direct electron detector (Gatan, USA) at the Harvard Cryo-Electron Microscopy Center for Structural Biology. Movie stacks were automatically recorded using AutoEMation in the super-resolution mode at a nominal magnification of 105,000 x, corresponding to a physical pixel size of 0.83 Å. The defocus was set to from −1.0 to −2.5 µm. A total exposure dose of 53 e^−^/Å^2^ was fractionated into 50 frames for each movie stack. We obtained one cryo-EM dataset of NiV L–P complex including a total number of 7803 movie stacks.

### Cryo-EM image processing

We processed all images using Relion 3.1^51^, movie frames gain-normalized and motion-corrected using MotionCor2^55^ and contrast transfer function (CTF) correction performed using CTFFind4.1^56^, as implemented in Relion 3.1. Particle-picking was performed in crYOLO ^53^ and 1,871,392 particles from 7,803 micrographs were picked. Picked particles were imported to Relion, extracted, binned to a pixel size of 1.66 Å and subjected to 2D classification. Good classes of particles from the 2D classification were selected and imported to CryoSPARC^52^ to generate an initial model. Three rounds of 3D classification with C1 symmetry were imposed on 1,842,231 particles, 477,936 particles of which were selected and subjected to auto-refinement to generate a 2.7 Å map. CTF refinement and Bayesian polishing were performed to improve the resolution. Consequently, a final map was generated to 2.3 Å. To improve the local resolution of P ⍺-helical cap structure, focused 3D auto refinement was performed, allowing us to obtain a masked a 3.0 Å map. We also used DeepEMhancer^57^ for cryo-EM volume post-processing.

### Model building and refinement

The structures of the five domains (RdRp, CAP, CD, MTase domain and CTD domain) of L and of monomeric P were predicted Phyre2^58^. These predicted structures and the crystal structure of the NiV P OD (PDB code: 4N5B, AAs 476–576) were used to build the atomic model of NiV L–P complex. The RdRp and CAP domains of L and tetrameric P OD plus a single P XD were docked and rigid-body fitted well into the cryo-EM map using UCSF Chimera. Extra residues of P were built manually in *Coot*^54^. We performed manual model building to improve local fit using *Coo*t^54^ and real space refinement using Phenix^50^. MolProbity was used to validate the geometries of the final models and the statistics are given in Table S1.

### 3D variability analysis

3D variability analysis (3DVA) was performed using cryoSPARC^40,52^. Particle stacks and reference map from Relion final 3D auto-refine were imported. 3DVA were performed using a mask generated from Relion 3.1. Twenty components were generated, of which major component showed flexibility in the P segment. The frames from these components were visualized and recorded in UCSF Chimera ^48^ as a volume series (see Movie S1).

### In silico modeling and molecular dynamics simulations

The cryo-EM structure of the NiV L–P complex was prepared before modelling and simulations. The module of Protein Preparation in Schrödinger Maestro^59^ was applied to cap termini, repair residues, optimize H-bond assignments, and run restrained minimizations following default settings.

The Schrödinger Desmond MD engine^60^ was used for simulations as previously described^61^. An orthorhombic water box was applied to bury prepared protein systems with a minimum distance of 10 Å to the edges from the protein. Water molecules were described using the SPC model. Na^+^ ions were placed to neutralize the total net charge. All simulations were performed following the OPLS4 force field^62^. The ensemble class of NPT was selected with the simulation temperature set to 300K (Nose-Hoover chain) and the pressure set to 1.01325 bar (Martyna-Tobias-Klein). A set of default minimization steps pre-defined in the Desmond protocol was adopted to relax the MD system. The simulation time was set to 100 ns for the protein system with three duplicate MD runs. One frame was recorded per 200 ps during the sampling phase. Post-simulation analysis of the RMSF was performed using a Schrödinger simulation interaction diagram. RMSF values from the Cα of each residue were used for plotting.

### Design and cloning of the NiV minigenome system

The minigenome system was designed based on sequences from the NiV Bangladesh strain (GenBank: AY988601.1). The cis-acting elements inserted into the multi-cycle replication minigenome were determined based on a previously described NiV minigenome template ^9^. The multi-cycle replication minigenome contained in 3’ to 5’ order, the first 112 nt of the NiV genome sequence, including the 55 nt *leader* region, *N gene start* signal, and 46 nt of non-translated sequence from the 3’ end of the *N* gene, which includes promoter element 2 (CNNNNN)_3_, 519 nt derived from the bacterial *chloramphenicol acetyltransferase* (*CAT*) gene, 105 nt of sequence derived from the *N-P* gene junction region (nucleotides 2247–2351 of the NiV Bangladesh complete genome) that includes the *N gene end* signal, trinucleotide intergenic, and *P gene start* signal, followed by 939 nt *Renilla luciferase* reporter gene sequence, and then the 5’ 100 nt of the NiV genome, including 56 nt of non-translated sequence from the 5’ end of the L gene, which contains the complement of promoter element 2, the L *gene end* signal and the 30 nt 5’ trailer region. Together with inserted restriction sites, the total minigenome length is 1794 nt. The minigenome cassette was flanked by a hepatitis delta virus ribozyme sequence adjacent to the *leader* region and a T7 promoter sequence adjacent to the *trailer* region (Figure S8A). This minigenome was used as a template to generate a single-step replication minigenome limited to the mini-antigenome step of replication. This minigenome had a deletion of promoter element 2 within the L nontranslated region and substitutions at position 1–4 relative to the 5’ end of the trailer region, which inactivated the trailer promoter at the 3’ end of the mini-antigenome (Fig. S8A). The multi-cycle replication minigenome followed the “rule of six” except for two additional G residues contributed by the T7 promoter; the single step replication minigenome followed the “rule of six” including the additional G residues contributed by the T7 promoter. Codon optimized versions of NiV Bangladesh strain N, P, and L open reading frames (Synbio) were inserted into a modified version of plasmid pTM1, which contains a T7 promoter and internal ribosome entry site. Each open reading frame was inserted such that the initiating codon was inserted into the plasmid NcoI site, and the 3’ end of the open reading frame was flanked with a 17 nt poly A sequence ^37^. In addition, the open reading frame of L contained a C-terminal strep tag. All plasmids were sequenced in their entirety (Plasmidsaurus) and their integrity and purity was confirmed by agarose gel electrophoresis prior to each transfection.

### Reconstitution of minigenome replication and transcription

BSR-T7 cells (a kind gift from Dr. Klaus Conzelmann)^41^ in six-well dishes were transfected with (per well) 0.3 µg of the relevant minigenome plasmid, 0.4 µg of N, 0.2 µg of P, 0.1 µg of L and 0.04 µg of firefly expression minigenome using Lipofectamine 3000 (Invitrogen). Each transfection reaction was set up in duplicate. Followed by incubation at 37°C for 22–24 h, the transfection mixture was replaced with OptiMem containing 2% (v/v) fetal bovine serum. Cells were harvested 44–48 h after transfection.

### Harvesting of transfected cells for Western blot and luciferase assays

For Western blot and luciferase analyses, cells from one transfected well were lysed in 500 µl passive lysis buffer (Promega). A 100 µl aliquot of the lysate was passed through a Qiashredder column (Qiagen) and a 14 µl aliquot was subjected to electrophoresis on an 8% SDS-polyacrylamide gel alongside a PageRuler Plus Prestained Protein Ladder (ThermoScientific). Proteins were transferred to nitrocellulose by Western blotting and the blots were probed with beta-tubulin rabbit polyclonal antibody (LI-COR) at 1:1000 and a Strep-tag classic mouse monoclonal antibody (Biorad) at 1:500. The primary antibodies were detected with donkey anti-rabbit and goat anti-mouse antibodies, each at 1:20,000. For luciferase assays, aliquots of cell lysates were diluted 1:30 and assessed for firefly and Renilla luciferase activity using a Dual-Luciferase Reporter Assay System (Promega) according to the manufacturer’s instructions. Quantification of the relative luciferase activity for each sample was performed by normalizing the Renilla luciferase value to the firefly luciferase value. The −L value was subtracted to account for background. The resulting values were then normalized to the corresponding value from the L wt reaction, which was set to 1.

### Northern blot and hybridization

For RNA analysis, cells from a transfected well were used for RNA extraction using a Monarch total RNA miniprep kit (NEB). Isolated RNA was subjected to denaturing gel electrophoresis alongside a molecular weight ladder (Dynamarker prestain marker for RNA high) in a 1.5% agarose gel containing 0.44M formaldehyde in MOPS buffer. The gel was transferred to a nylon membrane (Cytiva), which was stained with methylene blue to confirm that equivalent amounts of RNA were loaded in each lane and to allow visualization of the molecular weight markers. Negative- and positive-sense ^32^P-labled CAT-specific riboprobes corresponding to minigenome (or complementary) sequence wwere synthesized by T7 RNA polymerase. Following phenol-chloroform extraction, the appropriate riboprobe was hybridized to the membrane in 6X SSC, 2X Denhardt’s solution, 0.1% SDS, and 100 µg of sheared DNA per ml for a minimum 18 h at 65°C. The membranes were washed at 65°C in 2X SSC-0.1% SDS for 2 h and in 0.1X SSC-0.1% SDS for 15 min. Phosphorimager analysis was performed using an Azure Sapphire Biomolecular Imager and signals were quantified using Image Studio (LI-COR). A box of equivalent size as used for other bands of the same RNA species was drawn at the corresponding position on the −L sample lane and used to set the background to 0. The resulting values were then normalized to the corresponding value from the L wt reaction, which was set to 1.

### Figure preparation

UCSF Chimera and Pymol (Schrödinger LLC) were used for structure visualization and figure generation.

## Contributions

S.H. cloned, expressed, and purified recombinant NiV L–P complexes from insect cells for biochemical and structural analyses (negative stain and cryo-EM) and performed mass photometry experiments. S.H. also generated the initial model and built and refined the final model. H.K. generated the plasmids and optimized the protocols for the NiV multi-cycle and single-step minigenome systems. H. K. also performed transfections, luciferase, Western blot, and Northern blot analyses. P.Y. generated recombinant baculoviruses, purified recombinant proteins, vitrified samples for cryo-EM, collected and processed data, and generated 3D reconstructions of the NiV L–P complex. S.H., P.Y., Z.Y. participated in model building and refinement. P.Y. additionally conducted 3D variability analysis. B.L. performed in vitro RNA synthesis assays and designed and participated in cloning the L mutants. S.M. performed site-directed mutagenesis to generate L mutants and performed Western blot analyses. J.P. design and cloned L–P expression constructs. Y. B. performed molecular dynamics simulations and analyzed molecular dynamics data. S.H., H.K., P.Y., Y.B., R. F. and J.A. generated figures and participated in writing and editing the manuscript. M.S. participated in experimental design and analysis. R.F., J.A., and M.S. participated in funding acquisition. All authors contributed to discussions and reviewing the manuscript.

## Acknowledgements

We would like to thank the staff at the Harvard cryo-EM center (R. Walsh, R. Nair, C. Leistner, and M. Mayer) for help with data acquisition, pre-processing, and storage. This work was supported by an award from the Bill & Melinda Gates Foundation [INV-040438] to R.F.

